# High segregation and diminished global integration in large-scale brain functional networks enhances the perceptual binding of cross-modal stimuli

**DOI:** 10.1101/2020.09.02.276071

**Authors:** Soibam Shyamchand Singh, Abhishek Mukherjee, Partha Raghunathan, Dipanjan Ray, Arpan Banerjee

**Affiliations:** National Brain Research Centre, NH8, Manesar, Gurgaon 122052, India; Department of Psychology, Ashoka University, Sonepat 131029, India

**Keywords:** multisensory, whole-brain functional connectivity, pSTS, fMRI, segregation, integration, homeostasis, perception, predictive coding, Bayesian

## Abstract

Speech perception requires the binding of spatiotemporally disjoint auditory–visual cues. The corresponding brain network-level information processing can be characterized by two complementary mechanisms: functional segregation which refers to the localization of processing in either isolated or distributed modules across the brain, and integration which pertains to cooperation among relevant functional modules. Here, we demonstrate using fMRI recordings that subjective perceptual experience of multisensory speech stimuli, real and illusory, are represented in differential states of segregation–integration. We controlled the inter-subject variability of illusory/cross-modal perception parametrically, by introducing temporal lags in the incongruent auditory–visual articulations of speech sounds within the McGurk paradigm. The states of segregation–integration balance were captured using two alternative computational approaches. First, the module responsible for cross-modal binding of sensory signals defined as the perceptual binding network (PBN) was identified using standardized parametric statistical approaches and their temporal correlations with all other brain areas were computed. With increasing illusory perception, the majority of the nodes of PBN showed decreased cooperation with the rest of the brain, reflecting states of high segregation but reduced global integration. Second, using graph theoretic measures, the altered patterns of segregation–integration were cross-validated.

## Introduction

The illusion of a puppet vocalizing human speech that in reality comes from the sounds of a ventriloquist has intrigued mankind for ages (Alais and Burr, 2004). While spatial incongruency is a well-studied aspect of ventriloquism, perceptual binding of temporally dissonant stimuli is subtle (Bertelson and Aschersleben, 2003), such as when viewing a poorly dubbed video our perceptual system immediately perceives the asynchrony in sound and lip movements, without even understanding the language. Previous studies have also reported that within a certain window, the temporal dissonance between multiple sensory components can still facilitate illusory perceptual experience (Bertelson and Aschersleben, 2003; Munhall et al., 1996; Shams et al., 2002; Stevenson et al., 2012); but whether large-scale brain networks exhibit specific patterns that can explain the degree of crossmodal integration during perceptual binding is an unanswered question.

Several studies have highlighted that studying the interplay of segregative and integrative mechanisms offer a comprehensive theoretical framework to interpret large-scale distributed brain activations that are often reported in complex cognitive tasks (Cohen and D’Esposito, 2016; McIntosh, 2004; Mohr et al., 2016; Tomasi et al., 2014; Tononi et al., 1994). In this article, we hypothesized that the cooperation among relevant brain areas via functional integration mechanisms (Friston et al., 1993) and their segregative local information processing work harmoniously to facilitate the crossmodal perceptual binding, and this relationship may be measured using network neuroscience metrics and be driven by behavioral contexts which we aim to control parametrically. Previous studies have used the paradigms of sound-flash illusion (Watkins et al., 2007) and McGurk illusion (McGurk and Macdonald, 1976) to suggest posterior superior temporal sulcus (pSTS) as the key hub for crossmodal percept binding (Beauchamp et al., 2004; Nath and Beauchamp, 2011), whereas other studies have not found enhanced pSTS activation toward temporal synchrony of auditory and visual modalities (Jones and Callan, 2003). From a large-scale network perspective, noisy stimulus in one modality also increases functional connectivity between pSTS and areas of other modalities; e.g., a blurry video will enhance pSTS–auditory connectivity (Nath and Beauchamp, 2011). Such shifts in connectivity arising from stimulus associations can be linked to causal inference models of perception (Koerding et al., 2007; Magnotti et al., 2018), and lately, biophysically realistic network mechanisms have been proposed (Cuppini et al., 2017; Kumar et al., 2020; Thakur et al., 2016). Audio-visual speech perception also involves active participation of the motor network (Dehaene-Lambertz et al., 2006; Liberman and Whalen, 2000; Morís Fernández et al., 2017; Scott et al., 2009), and during the presentation of incongruent (McGurk) speech stimuli, there is increased interaction between motor representational area and inferior frontal gyrus. The motor network was however identified to be contributing towards resolving the audio-visual conflicts (Murakami et al., 2018). The manifestation of audio-visual illusory speech perception thus involves functional domains promoting the integration, such as the pSTS network, and domains with more abstract functions, such as the motor network. The present study empirically demonstrates how subjective perceptual binding emerges from synergistic functional integration and segregation mechanisms of brain (Tomasi et al., 2014). Moreover, the existence of an altered balance in network mechanisms can explain why perception is relatively stable for a wide range of behavioral contexts (Zopf Jr, 1963), e.g., perception of illusory sound for different vocalizers and post-learning (Mallick et al., 2015), perceivers across cross-cultural boundaries (Magnotti et al., 2015), even though inter-individual variability exists.

To elicit instances of cross-modal perceptual experience, the McGurk incongruent video—auditory /pa/ superimposed on the lip movement of /ka/ from a human vocalizer (McGurk and Macdonald, 1976)—was shown to volunteers during an fMRI experiment (Fig. 1A). Subsequently, subjective perception was parametrically controlled by using different versions of this video created by introducing temporal lags between the auditory and visual sensory streams. The temporal lag ranges from −300 (A leading V) to 450 ms (V leading A) in intervals of 150 ms (Fig. 1A); also shown are matched number of synchronous congruent /ta/ (auditory /ta/–visual /ta/). The choosing of an asymmetrical range of audio-visual lags (−L2, …, L0, …, L3) was guided by previously reported perceptual bias toward visual-leading scenarios (Munhall et al., 1996). Volunteers reported whether they heard /ta/, /pa/, or “others” in a forced-choice paradigm.

**Figure 1.**
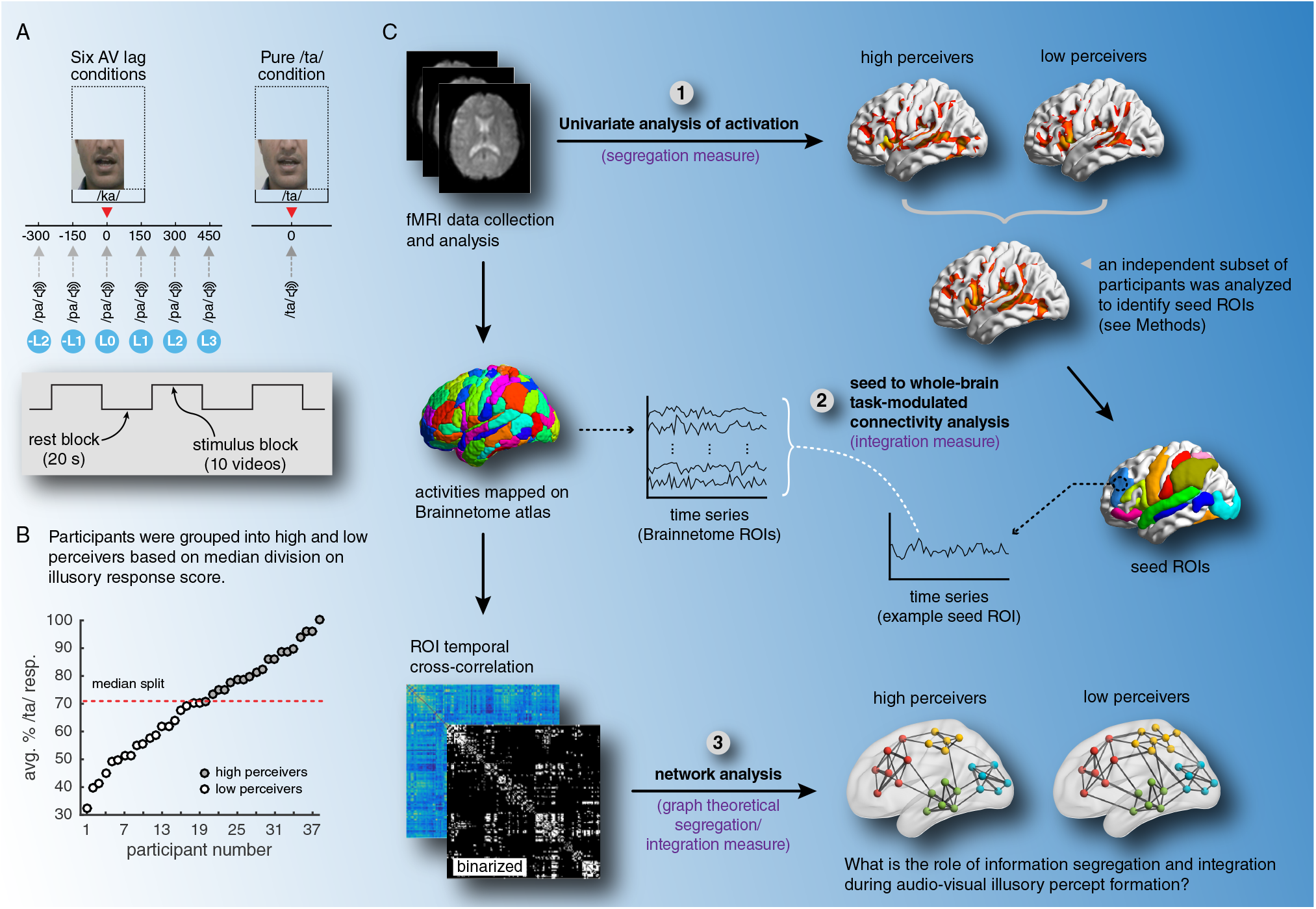
(*A*) Stimulus design using the McGurk paradigm. (*B*) Intersubject variability in susceptibility for illusory perception for all 38 selected volunteers; the susceptibility was evaluated as the average of two maximal percentage /ta/ responses in any two AV lags (dotted line indicates the division between high and low perceivers). (*C*) Schematic showing various analysis steps used to study the integration/segregation balance.

Cognitive tasks generally involve distributed brain areas, and a riveting hypothesis for normative human cognition is the maintenance of homeostatic balance between integrative and segregative mechanisms (Cohen and D’Esposito, 2016; McIntosh, 2004; Mohr et al., 2016; Tomasi et al., 2014; Tononi et al., 1994), with imbalances leading to neuropsychiatric disorders (Lord et al., 2017). The present study also demonstrates how alterations of balance affects the perceptual binding of incongruent multisensory stimuli. This is a significant departure from the earlier understanding of multisensory integration from the perspective of inverse effectiveness which has been reinforced by several studies using different neuroimaging modalities (Meredith and Stein, 1983; Molholm et al., 2002; Stevenson and James, 2009).

## Results

### Parametric Control of Illusory Perception

For the McGurk incongruent stimulus, more than a chance level of varied responses to illusory cross-modal /ta/ perception were observed in all participants at multiple AV lags, demonstrating that indeed the degree of AV temporal synchrony affects subjective perception (Fig. 2 and Supplementary Fig. 1). In order to study the neural basis of subjective variability, we categorized the participants into two equally sized groups (*N* = 19) after being sorted based on their *susceptibility* to audio-visual illusion. The susceptibility was measured as the average of two maximal percentage /ta/ responses in any two AV lags. The group reporting higher illusory perception is termed as *high perceivers* and the other group at the lower end as *low perceivers* (Fig. 1B).

**Figure 2.**
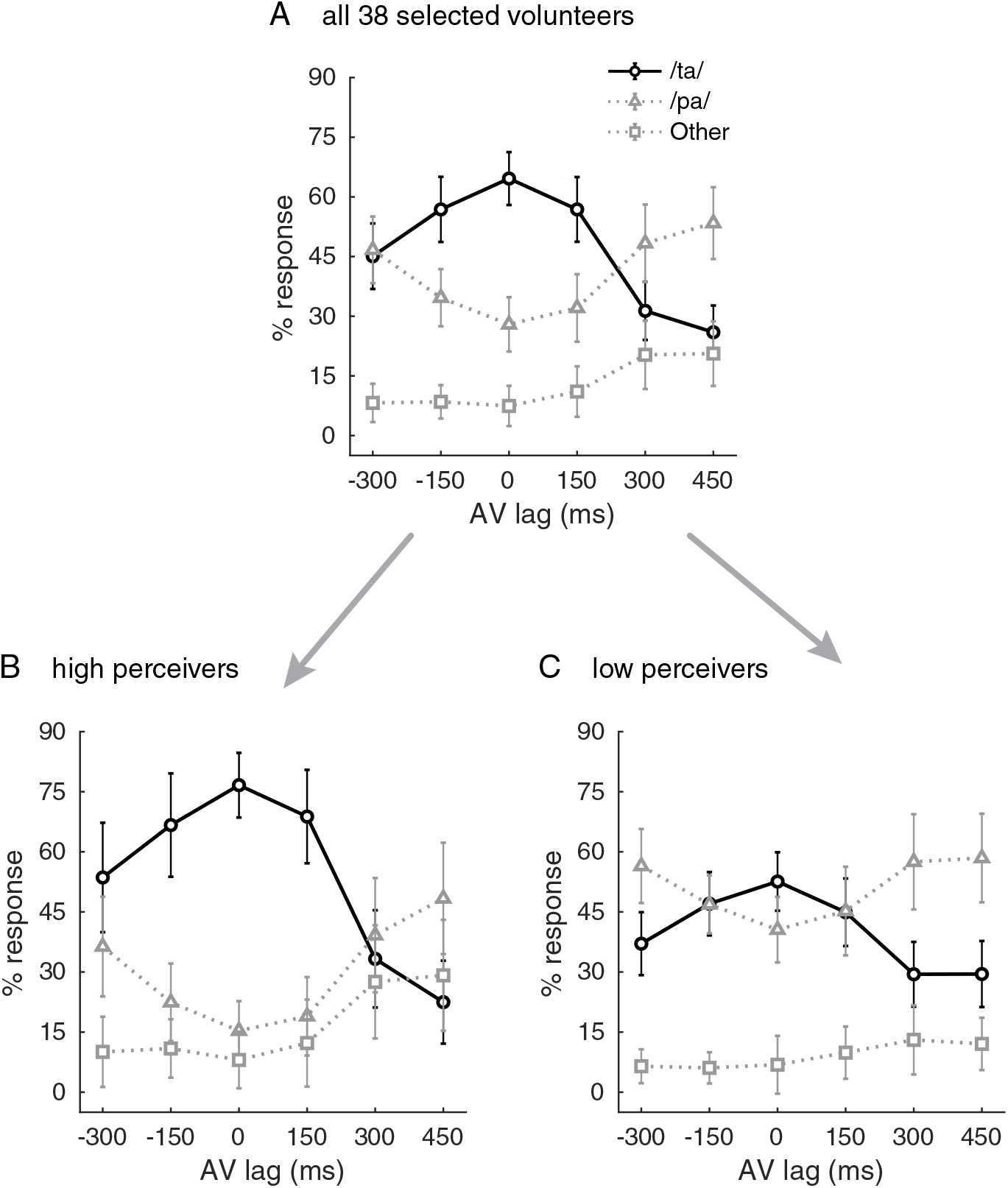
(*A*) Behavioral responses averaged across all 38 selected volunteers. Behavioral responses averaged across (*B*) high perceivers (*N* = 19) and (*C*) low perceivers (*N* = 19). Error bars represent the standard error of the mean (95% CI) across the respective population.

A two-way ANOVA was further conducted to reveal the effect of AV lag and subject grouping factors on participants’ perceptual responses. In the case of illusory /ta/ responses, there were significant main effects of AV lag (*F* (5, 213) = 17.9202, *p* < 0.0001), as well as of participants’ grouping (*F* (1, 213) = 20.1903, *p* < 0.0001) at 95% confidence levels (a confidence interval used in all statistical significance tests in the article). We also observed a statistically significant interaction between the two factors (*F* (5, 213) = 2.9257, *p* = 0.0141); Tukey’s post-hoc comparison further showed that, between the same time lag condition, high perceivers were more susceptible to illusory /ta/ perception than the low perceivers at L0 (*p* = 0.0459) and L1 (*p* = 0.0493) conditions. For the unisensory /pa/ response, there was no significant interaction between the two factors (*p* = 0.7087). However, significant main effects of AV lags (*F* (5, 213) = 6.6862, *p* < 0.0001), as well as of subject grouping (*F* (1, 213) = 40.7071, *p* < 0.0001) were observed. Similarly, no significant interaction between the two factors was observed while reporting “other” responses also (*p* = 0.3774), but we again observed significant main effects of AV lags (*F* (5, 213) = 3.5001, *p* = 0.0046) and participant grouping (*F* (1, 213) = 7.3264, *p* = 0.0073).

### Brain Mapping of Perceptual Binding

fMRI images were analyzed using SPM software (Friston et al., 1995) (see Fig. 1C for the schematic of the analysis pipeline). Group-level inference from random effect analysis revealed that maximal brain areas were active during L0 and L1 conditions, where propensities to cross-modal illusory perception were also maximally reported (Fig. 3A). The extent of task-activated brain areas diminished bilaterally with increasing AV separation (Fig. 3A). Furthermore, when compared with high perceivers, the low perceivers showed relatively weaker activation especially in primary motor, right frontal, anterior cingulate cortex, superior temporal region including pSTS, rostral inferior parietal, and occipital areas; on the other hand, orbitofrontal, insula, and thalamic regions showed higher activation in low perceivers.

**Figure 3.**
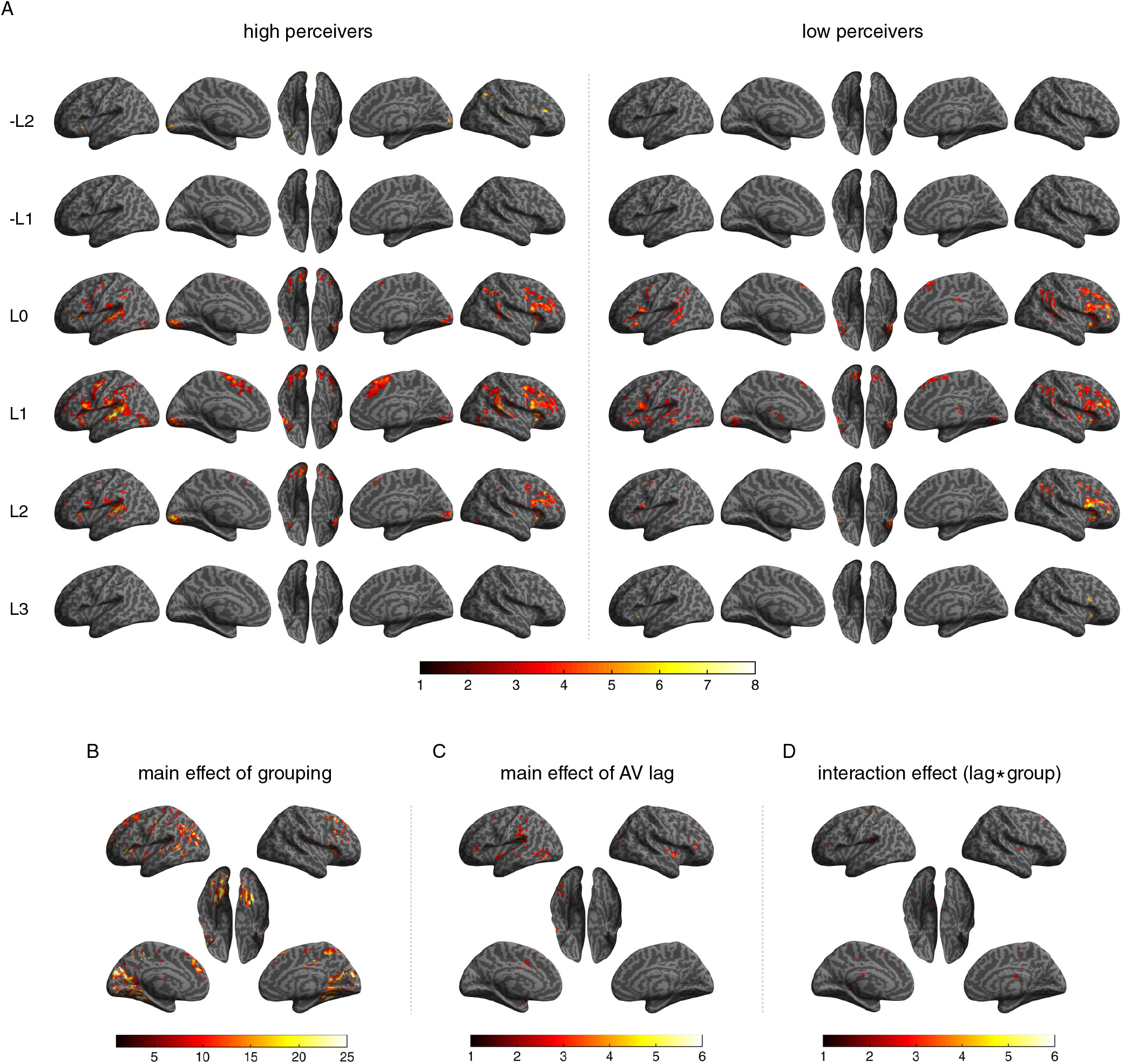
(*A*) Group-level task-enhanced (task > rest) active brain areas for high and low perceivers (peak *p* ⩽ 0.05, FDR corrected). Color bar represents the T-score. (*B*) Main effect of participants’ grouping factor. (*C*) Main effect of AV lag factor. (*D*) The interaction between the groupings and AV lag. Color bars in panels (*B–D*) represent the F-score. The activations during congruent /ta/ stimuli are reported in Supplementary Figure 2.

We further performed a full factorial analysis of the activations with time lag and participants’ grouping as independent factors. The analysis was done in SPM with the default settings. We observed significant main effects of participants’ grouping (*F* (1, 213) = 7.3491, *p* = 0.007) as well as for AV lag (*F* (5, 213) = 2.256, *p* = 0.0049) (Fig. 3 B and C). The grouping factor had a significant effect on clusters located in all the brain parcels that were found active during L0 and L1 conditions of high and low perceivers. Significant main effect of AV lags was observed in clusters located dominantly in inferior frontal gyrus, superior temporal gyrus including pSTS, caudal cingulate cortex, inferior parietal lobule, insula, lateral occipital cortex, and thalamus. Interestingly, a significant interaction between the two factors was also observed (*F* (1, 213) = 2.256, *p* < 0.0048), and the active clusters contributing to this interaction were found majorly in superior and inferior frontal gyri, precentral gyrus, inferior parietal lobule (rostroventral area 40), caudal cingulate cortex, and thalamus (Fig. 3D). The clusters with significant effects are reported in Supplementary Table 1, with the corresponding brain subregions mapped according to Brainnetome parcellation (http://atlas.brainnetome.org/; Supplementary Table 2).

To study the differential functional roles of associated brain regions during audio-visual crossmodal perception, 29 seed regions were identified from the group-level analysis of all the 38 participants during synchronous congruent Pure /ta/ condition (Fig. 1A; Supplementary Fig. 2A; Supplementary Table 3). We will henceforth term these 29 seed ROIs which were also found active during L0 and L1 conditions as the perceptual binding network (PBN). The rationale behind using the congruent Pure /ta/ condition for the seed ROI selection are: (1) variable illusory perceptual response from the participants (although the stimulus is congruent) reflects the underlying cross-modal sensory processes. The active brain regions will thus reflect the areas responsible for both unisensory and multisensory processes. (2) The congruent condition also helped us to avoid circularity in the seed ROI selection.

### Synergistic Network Organization

The process of cognition involves interactions among associated neural units (brain regions). To investigate the synergistic interactions involving the 29 seed regions of PBN, the seed-based whole-brain functional connectivity (SWFC) for each seed was computed through gPPI modeling (Fig. 1C; see also *Methods* for details) (Friston et al., 1997; McLaren et al., 2012). It was observed that the averaged SWFC mostly decreased at L0 or L1 for high perceivers (see the example left SFG seed regions, Fig. 4A), and the trend is either reversed or weakly followed in the case of low perceivers (Fig. 4B). The question now is whether this decrease in SWFC can be meaningfully associated with the propensity for illusory perception. To meet this end, SWFC– behavior correlation (SBC; see *Methods*) was computed for each PBN seed region. We observed that for high perceivers the majority of the seed ROIs displayed negative SBC, with the positively correlated seed regions mostly concentrated in frontal brain areas and in the right hemisphere (see Fig. 4C and Supplementary Table 4). On the other hand, the low perceivers displayed either a positive or a weakly negative SBC values. These findings emphasized that globally decreased information integration through the PBN seed regions was essential for the illusory audio-visual percept formation. Furthermore, to investigate if the results are independent of the choices of any particular brain parcellation scheme, the procedures were repeated using Schaefer atlas having 300 ROIs (https://github.com/ThomasYeoLab/CBIG/tree/master/stable_projects/brain_parcellation/Schaefer2018_LocalGlobal/Parcellations/MNI) (Lawrence et al., 2021; Schaefer et al., 2018). We observed a similar pattern of decreasing global information integration from the seed regions with increasing audiovisual illusory perception (Supplementary Fig. 3).

**Figure 4.**
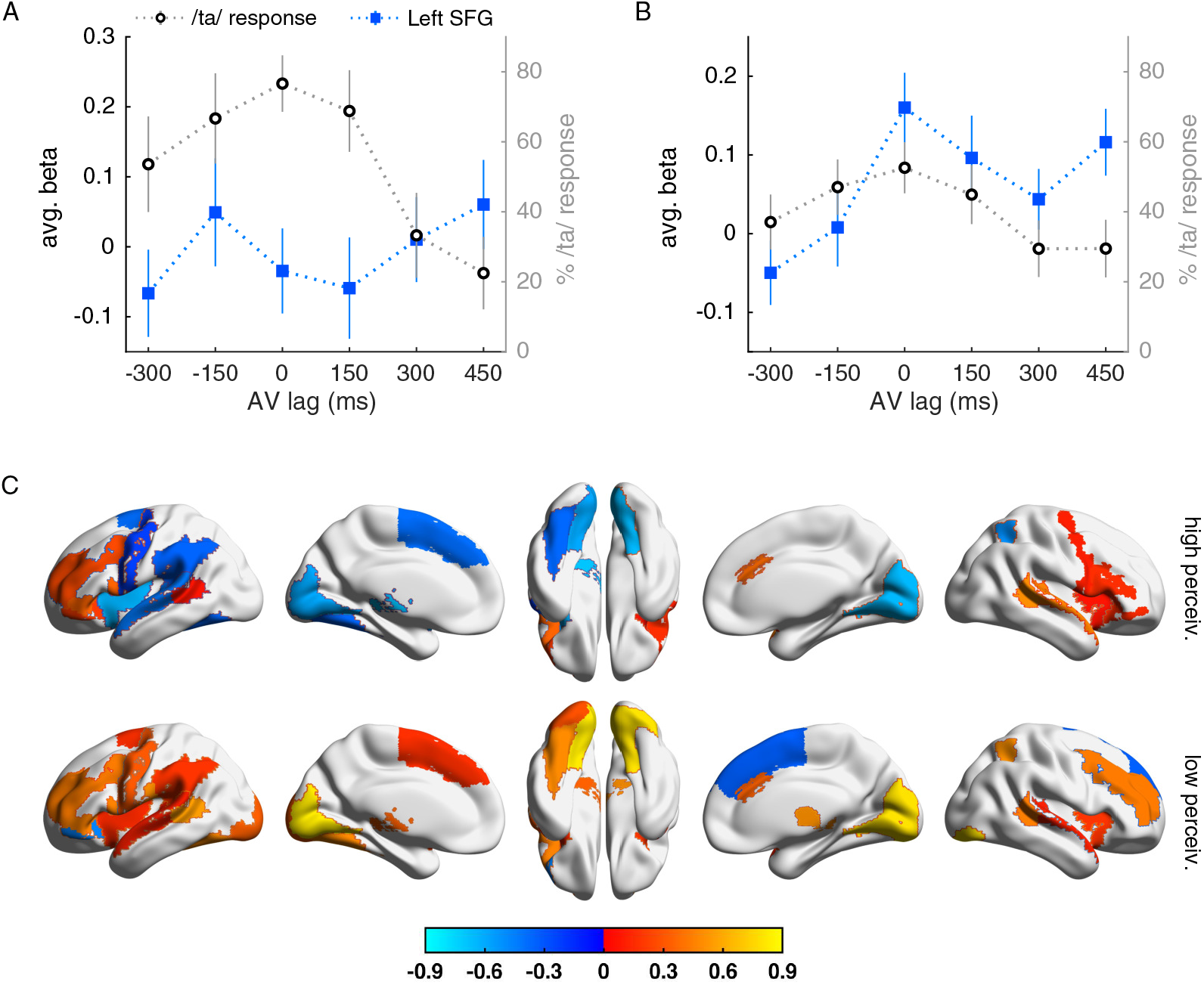
(*A*) Average seed-based whole-brain functional connectivity (SWFC) changes across AV lag conditions for an example left SFG seed in the case of high perceivers, compared against their degree of illusory /ta/ perception (black dotted line). (**B**) The corresponding plot for the low perceivers. Error bars represent the standard error of the mean (95% CI). (**C**) Surface plot showing the strength of SWFC–behavior correlation (SBC) for all 29 seed ROIs of PBN in high perceivers (upper) and low perceivers (lower) (cf. Supplementary Table 4; see also *Methods* for details).

To validate the above finding that hints toward an altered information segregation–integration balance during the cross-modal perceptual binding, we conducted an independent investigation on the topological features of whole-brain functional networks using metrics from graph theory. For this analysis, 50 binarized whole-brain network subsamples were constructed at the group level for each stimulus condition by randomly permutating 70% of the participants in each group—high and low perceivers. For estimating the network segregation, averaged local clustering coefficient and modularity measures were used; the integration measures include the characteristic path length and averaged node participation coefficient. Each network measure was further normalized with estimated measures from randomized networks (see *Methods* for more details). In the case of high perceivers, the normalized clustering coefficient (*C*_*n*_) increased with decreasing AV lag and peaked at L0 where the reported average propensity to illusory perception is also the highest (Fig. 5A). This indicates there is increased local cohesiveness in the network with increasing illusory perception. However, contrary to this effect is the increase in normalized characteristic path length (*L*_*n*_), indicating a decrease in global cohesiveness. The characteristic path length may be defined as the minimum steps each node pair will take to reach each other, averaged across all the available node pairs in the network. So, it does not however include the factor of actual structural connection lengths between brain regions. Hence, the manifestation of audio-visual illusion might thus require an altered balance between the segregative (local cohesion) and integrative (global cohesion) information processing. Increased illusory experience was also associated with increasing dissociation among the functional modules, as reflected from the increased normalized modularity score (*Q*_*n*_). The functional dissociation can also be interpreted from the decrease in average node participation coefficient (*P*) with increasing illusory perception (Fig. 5A). Thus, the altered balance between functional segregation and integration can be characterized from the node-level measure (e.g., local clustering coefficient, participation coefficient), and also from the whole-network-level measure (e.g., normalized modularity score). Low perceivers also displayed a similar trend, however the maximum values of clustering coefficient, characteristic path length, and modularity, and also the minimum value of participation coefficient were observed in L1 condition. The above results were also stable against the choice of a different parcellation scheme (Supplementary Fig. 4).

**Figure 5.**
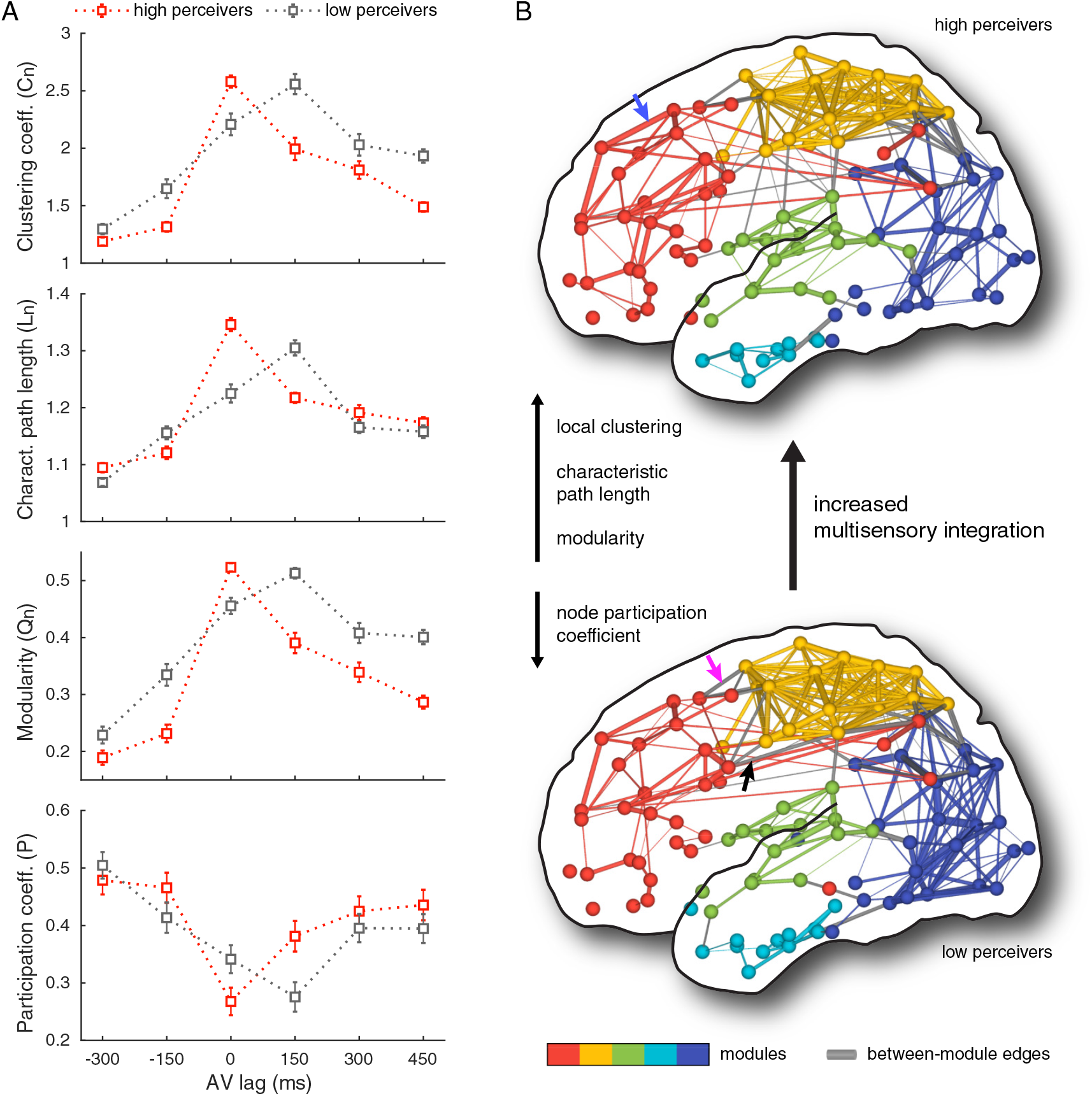
(*A*) Change in the network topological measures for high and low perceivers. (*B*) Left hemispheric brain network differences between high and low perceivers during L0 (synchronous and incongruent) condition.

Taking the left hemispheric brain networks at L0 condition as an example, the network differences between high and low perceivers are illustrated in Fig. 5B: e.g., the modules in the frontal and temporal brain regions show increased within-module connectivity both in number and strength (cf. blue arrow), and decrease in-between-module connectivity (cf. magenta and black arrows) in the case of high perceivers. However, other AV lag conditions did not necessarily show similar trends. One possible explanation for this inconsistency is the difference in the activation patterns between the two groups, and also the decreasing effect of illusory perception with increasing AV lag. The reduced activation was specifically observed in many sensory as well as associative cortical regions of low perceivers, and this difference in activation pattern might shift the segregation–integration balance accordingly (Fig. 3A). The graph theoretic measures thus delivered a similar take-home message, as observed for SWFC with behavior, that increased cross-modal illusory perception triggers increased local segregation realized via an increase in clustering coefficient, characteristic path length, and modularity; and also the minimum participation coefficient.

## Discussion

We demonstrated that patterned relationships between segregative and integrative information processing mechanisms employed by distributed brain areas are critical for the binding of auditory (A) and visual (V) signals into a coherent perceptual experience. A distributed ensemble of brain regions defined as the perceptual binding network (PBN) exhibited heightened activity when A and V were either temporally congruent (L0) or nearly congruent (L1) (cf. Fig. 3A). The heightened activity in PBN nodes can be interpreted as local segregation whereas the decreased functional connectivity with the rest of the brain at L0 and L1 reflects diminished global integration. Hence, the simultaneous occurrence of both increased local segregation and diminished global integration can be defined as an altered state of balance between integration and segregation mechanisms that results in higher propensity to perceive illusory-McGurk (cross-modal) sounds. When the level of binding traverses from low to high states, the pattern of balance between segregation and integration mechanisms undergoes an alteration. Supporting this emerging view, we also observed that Pure /ta/ activations in low perceivers (Supplementary Fig. 2B) were similar to the incongruent McGurk stimuli activations at L0/L1 whereas activation levels in high perceivers for Pure /ta/ were overall very low. The corresponding states of altered balanced can also be characterized exclusively from graph theoretic network measures. Finally, the increased thalamic activity in low perceivers also supported an enhanced role of subcortical level processing during cross-modal perception (Gau et al., 2020; Noesselt et al., 2010; Tyll et al., 2011).

Previous studies focused on the participation of individual regions such as pSTS for multisensory processing and integration (Nath and Beauchamp, 2011; Stevenson and James, 2009). However, several unanswered questions related to multisensory integration and more subtly cross-modal binding (how stimulation of one sensory stream affects others) require a theoretical understanding of distributed network mechanisms (Johnston et al., 2022; Yau et al., 2015). Our paradigm allowed us to distinguish multisensory processing (involved in all AV lags, as well as during congruent AV stimuli) from the cross-modal perceptual binding (predominantly in L0 and L1), and the importance of the distributed brain regions which together was defined as PBN. Confirming previous studies (Beauchamp, 2005; Beauchamp et al., 2004), our data clearly establishes pSTS activation is enhanced during crossmodal illusory /ta/ perception (Fig. 3A). However, as was also observed in L0 and L1 stimulus conditions where the degree of pSTS activation could not solely explain the perceptual difference between high and low perceivers, the interaction among distributed brain areas encompassing the primary sensory areas, motor representational areas, associative areas such as IPL, V5/MT+, IFG, etc., is essential for understanding audio-visual illusory percept formation. Hence, departing from the earlier studies, our results underscore the need for building an understanding of cross-modal perception at the whole-brain network level. Interestingly, we found more thalamic activation in low perceivers which indicated a major role played by the cortico-thalamic circuitry in perceptual binding. Recent studies have demonstrated that a decrease in levels of alpha coherence in electroencephalographic (EEG) activity is indicative of the propensity to McGurk-illusion perception (Kumar et al., 2020), wherein the cortico-thalamic basis of alpha rhythms is well-known (da Silva et al., 1973) as well as cortico-thalamic influences on early sensory cortices (Noesselt et al., 2010). fMRI time-scale may not allow the temporal resolution to identify the dynamic mechanisms of the cortico-thalamic circuitry, but future electrophysiological studies can surely address this question.

The second important finding of this study was the decrease in global integration of information effectuated by PBN nodes with the rest of the brain with increasing propensity to cross-modal perceptual experience. This was captured via the decrease in task-induced function dependence among several task-activated ROIs and the rest of the brain areas as a function of increasing cross-modal perceptual experience (Fig. 4). From the whole-brain network perspective also, increased cross-modal binding is associated with decreased global integration simultaneously co-occurring with increased local cohesiveness. We argue that the competitive balance of integrative and segregative processing delimits an individual’s perceptual experience, and alteration of the patterns defined by the individual’s functional connectome can explain perceptual variability (Finn et al., 2015). To the best of our knowledge, this is the first demonstration of competitive segregation and integration mechanisms in multisensory integration tasks. We thus argue that integrative mechanisms are predominantly compensatory compared to the task-driven haemodynamic oxygen metabolism, while contextualizing our results with previous work on multisensory integration (Nath and Beauchamp, 2011). Temporal binding windows (Stevenson et al., 2012) can define the boundaries of the compensatory regimes where integration and segregation can be synergistically altered, giving rise to inter-trial variability.

Our study departs significantly from the earlier theoretical frameworks in providing a more neurobiologically grounded mechanism of multisensory integration based on the existence of balanced states of higher segregation and decreased global integration (Lord et al., 2017) employed by nodes of the PBN. In light of our current results and previous modeling studies (Abeysuriya et al., 2018; Naskar et al., 2021; Vattikonda et al., 2016), we propose that balanced states of functional integration and segregation may emerge from the underlying excitatory–inhibitory (E-I) balance in local neuronal populations of involved brain areas and may even be related to neurotransmitter homeostasis (Naskar et al., 2021; Shine et al., 2021). While spontaneous brain activity such as during resting state has been typically associated with the E-I balance (Abeysuriya et al., 2018; Deco et al., 2017; Naskar et al., 2021; Vattikonda et al., 2016), the potential of altered E-I balance—in guiding the formation of perceptual experience from interactions between inclement sensory signals with stored associative memories (Barron et al., 2017), as a catalyst for volitional control effectuating perceptual stability (Kondo et al., 2018), in guiding decision-making strategies during multi-attribute choice (Pettine et al., 2021); all have been proposed recently. According to this line of reasoning, an altered balance between segregation and integration observed during the illusory perception may be the outcome of energy consumption by the motor networks as well, reported in an earlier study (Murakami et al., 2018) and supported by underlying neuromodulatory systems which are otherwise adapted to lesser energy consuming congruent auditory–visual input. The existence of the stable illusory perception across groups of people from wide geographical–linguistic backgrounds (Magnotti et al., 2015), as well as its robust presence post-learning (Mallick et al., 2015), also hints toward a neuromodulatory origin. An immediate goal of subsequent studies should be to employ neurostimulation techniques such as TMS and tDCS (Yau et al., 2015) with a goal to perturb the balanced states of integration–segregation mechanisms focusing on the PBN regions. An important limitation of the present study was that, with decreasing AV lag, although the graph theoretic measures such as characteristic path length that measures the functional segregation showed an increasing trend, simultaneously with a decreasing trend for participation coefficient that quantifies the functional integration, when these network measures are correlated with the degree of illusory perception (percentage /ta/ response) across all participants the trends were not significant even for the L0 condition. Hence, in future, a larger sample size will be required to evaluate the participant specific behavior. This brings us to another important limitation of the current study, that due to the block design which have inadequate repetitions of blocks, a subject-specific activation analysis and correlation with behavior couldn’t be undertaken. However, in future studies, an event related design augmented by collection of fMRI data at fewer lags will give us the statistical power to evaluate the finer reorganization of functional connectivity differences. Extending this line of reasoning, studies on schizophrenia and other schizoaffective disorders can explore whether an alteration of the balance of segregation and integration can be a viable framework to characterize patient-specific symptoms (Silverstein and Lai, 2021; Waters et al., 2014).

## MATERIALS AND METHODS

### Participants and Stimuli

Fifty-two healthy right-handed human volunteers (25 female, 27 male; mean age = 24.5 years, SD = 3.12) participated in the functional magnetic resonance imaging (fMRI) study. The study was approved by the Institutional Human Ethics Committee (IHEC), National Brain Research Centre, India, and all participants provided informed consent in a format approved by the IHEC.

Seven video stimuli each of 2 second duration were prepared. Six of them were incongruent audio-visual (AV) objects where audio recordings of a male speaker vocalizing /pa/ is dubbed on a video showing the lip movement while articulating the vocalization of /ka/ (/pa/–/ka/). The remaining one video was of a congruent AV object (/ta/–/ta/). In the six incongruent video stimuli, the gap between the onset of auditory and visual streams varied from −300 to 450 ms in steps of 150 ms (Fig. 1A). Negative sign indicates that the auditory stimulus onset precedes that of visual lip movement (A leading), and the positive sign on the other hand implies the auditory stimulus succeeds the lip articulation (V leading). An asymmetric range of AV lags was chosen because Munhall and colleagues (Munhall et al., 1996) reported the dominance of illusory perception toward positive lags where the onset of lip articulation precedes auditory onset. In the stimulus video, the articulation of /ka/ or /ta/ always started from a neutral lip-closed position. The congruent /ta/–/ta/ stimulus had synchronous onset of AV stimuli. Videos were created using VideoPad video-editing software (NCH Software, CO, USA) at a frame rate of 25 frames/second and a resolution of 1280 × 720 pixels. The auditory /pa/ and /ta/ syllables were of 0.549 and 0.531 s duration and were edited to minimize the background noise in Audacity software (Sourceforge.net). The audio sampling rate was 44 kHz and had a bit rate of 128 kbps. The resulting seven stimuli thus have five asynchronous and two synchronous stimuli— McGurk stimuli (/pa/–/ka/) with −300, −150, 0, 150, 300, 450 ms AV lags, and Pure /ta/ (/ta/–/ta/) stimulus with 0 ms AV lag. Henceforth, we shall also refer to the AV lag conditions in shorter notation as −L2, −L1, L0, L1, L2, L3 for −300, −150, 0, 150, 300, 450 ms lag, respectively.

### Task Design

The task design is illustrated in Fig. 1A. Inside the scanner, the stimuli were presented in a block design. Each activation block of length 20 s comprised 10 video stimuli of one catgory (L0/L1/…, etc.). In total there were 28 activation blocks in the whole experiment: 4 repeats for each of the 7 stimulus categories. The 28 activation blocks were alternated with another 28 inter-trial resting blocks or similar duration (i.e., 20 s) where participants attended to a fixation cross. We randomized the presentation of the 7 stimulus categories. The stimuli were presented through an INVIVO MR-compatible CRT screen attached to the head-coil and an MRI-compatible headphone (Philips, The Netherlands). Presentation software (Neurobehavioral Systems, CA) was used to display the stimuli. The participants were instructed to attend to the AV stimuli watching at the male speaker and report their perceptual experience of whether they heard “/pa/”, “/ta/,” or “any other” through a three-button fiber-optic response device by Curdes as soon as possible (Current Designs, PA, USA). They were also instructed that in case they missed a response, not to worry about the performance and rather concentrate on the current video being played. Most importantly, the volunteers were not given any cue on which AV lag condition will appear next. They were also not given any feedback on their response, since there is no “right” or “wrong” response in the incongruent conditions.

### fMRI Data Acquisition

Structural T1-weighted magnetization-prepared rapid gradient-echo (MP-RAGE) scan was performed at the beginning of each participant’s scanning session using a 3T Philips Achieva MRI scanner. The T1-weighted rapid images were acquired with TR=8.4 ms, TE=3.715 ms, and flip angle=8^°^; the field of view was 250×250×170 mm^3^ with a resolution of 0.977×0.977×1 mm^3^. For each participant, seven functional T2* scans (TR = 2000 ms, TE = 35 ms, flip angle = 90^°^) corresponding to the seven different stimuli (−L2 to L3 and Pure /ta/) were acquired. An 8-channel head coil was used to acquire the T2*-weighted images. The voxel size of the acquired images was 3.59×3.59×5 mm^3^ and the voxel size of the reconstructed images was 1.80×1.80×5 mm^3^. The slice thickness was 5 mm and inter-slice distance was 1 mm.

### Behavioral Data Analysis

The button press responses from the 52 participants were converted into normalized percentage measures of each perceptual response category—/pa/, /ta/, or “other”. A maximum of 40 responses were expected from each task. Tasks with less than 80% responses were rejected, however all participants met this threshold. The volunteers were further screened based on their /ta/ response percentage during congruent Pure /ta/ (/ta/–/ta/) stimulus condition. The minimum threshold of /ta/ response was set to 40%, and seven participants were rejected from this criterion. Two more participants were rejected because one had reported >95% /ta/ response in five AV lag conditions and another >95% “other” response in two AV lag conditions. Thus, on the basis of their behavioral responses we selected 43 participants. Five more participants had to be rejected because of excessive movement during MRI scanning. Finally, 38 participants were thus selected for further analyses. For each participant, the *susceptibility* to audio-visual illusion was estimated by averaging maximal percentage /ta/ responses at any two AV lags. A median split to their susceptibility scores allowed the participants to be binned into two equal-sized groups (19–19): the group representing higher susceptibilities will be called *high perceivers* and the other at the lower end as *low perceivers* (Fig. 1A). Statistical analyses of the perceptual responses—/pa/, /ta/, and “other”—were performed using two-way analysis of variance (ANOVA) with AV lags and participant grouping as independent variables.

### fMRI Data Analysis

#### Pre-processing

For each participant, the acquired functional brain volumes were realigned and co-registered with the structural scan using a six-parameter rigid-body transformation (three translations and three rotations). Those participants with maximum movement greater than 2 units (i.e., 2 mm in translational axes, or 2^°^ in rotational axes) in any of the six degrees of freedom were rejected from further analysis. The structural T1-weighted brain image was then segmented into grey and white matter maps, and the generated forward deformation field was used to spatially normalize the functional brain volumes to the Montréal Neurological Institute (MNI) standard brain employing the standard trilinear interpolation method. The images were resampled to 2×2×2 mm^3^ resolution. The normalized images were then spatially smoothed minimally with a 4 mm full-width at half-maximum (FWHM) isotropic Gaussian kernel. The preprocessing was done using Statistical Parametric Mapping Software (SPM) in MATLAB (MathWorks, MA). The first six brain volumes of each participant were removed to allow the magnetization to reach to a steady state.

#### Stimulus-based activation

After pre-processing, we performed the first-level fixed-effects analysis on the activation blocks versus the inter-trial rest blocks, with the experimental design (event onset and their time course) as essential regressors and the realignment parameters for head movements as nuisance regressors. Using the contrast images (task block > inter-trial rest block) from the first-level analysis, a second-level random-effects analysis was performed to generate group activation maps (Friston et al., 1995) for each AV lag condition, which captured the stimulus-driven changes. To study the main and possible interaction effects on the BOLD activations, a full factorial analysis was also performed with AV lags and participants’ grouping as independent factors (Supplementary Table 1).

The present study also identifies a set of seed regions of interest (ROIs) which together could be responsible for the audio-visual illusory percept formation. To define this set, we performed group-level activation analysis of the congruent synchronous Pure /ta/ condition (cf. Fig. 1A), inclusive of all the 38 participants. We used the Pure /ta/ stimulus condition after considering the fact that participants reported variable illusory perceptual experiences during this stimulus condition, reflecting an underlying cross-modal sensory integration process. The involved brain areas will thus represent the necessary unisensory and multisensory processing components. Another advantage of using the congruent condition is to avoid the circularity in ROI selection. These active brain parcels were also majorly found to be active during L0 and L1 conditions of high and low perceivers. The active parcels were then grouped into 29 ROIs based on major anatomical landmarks of Brainnetome atlas (Fan et al., 2016) (http://atlas.brainnetome.org/; Supplementary Table 2 and Table 3).

### Functional Connectivity Analysis

The 29 seed ROIs identified from SPM analysis were used to study functional integration mechanisms by computing the pairwise functional connectivities between each seed ROI and the rest of the brain (*seed-based functional connectivity*). While this approach identifies the global functional integration of a given seed, which when related to seed activation patterns identifies the segregation–integration balance; a more direct way to evaluate this key aspect can come from the *whole-brain graph theoretic network analysis*. Hence, we used both these approaches to study the balance of functional integration–segregation mechanisms.

#### Seed-based functional connectivity

For the seed-based connectivity analysis, each participant’s grey matter volume was parcellated using the Brainnetome atlas. Among the 210 cortical regions of the atlas, the two subregions of pSTS in each hemisphere were merged into a single ROI. From the subcortical regions, only the whole thalamic nuclei in each hemisphere were considered. Taken together, the final set of 210 ROIs covered the entire cortical region and the thalamus (Supplementary Table 2). To measure the task-modulated seed-based functional connectivity changes across AV lag conditions, the functional interactions between each 29 seed region and the remaining 210 ROIs (i.e., exclusive of seed parcels) were evaluated using the generalized psychophysiological interaction (gPPI) modeling approach (Friston et al., 1997; McLaren et al., 2012). We used the CONN toolbox to implement the gPPI model (Whitfield-Gabrieli and Nieto-Castanon, 2012). For each participant, the averaged beta value representing each seed’s functional connectivity during a particular AV lag condition was evaluated. We will term this measure as seed-based whole-brain functional connectivity (SWFC).

The relationship between the trend that SWFC changes across AV lag conditions and the corresponding changes in the degree of illusory /ta/ perception was further evaluated by estimating the Pearson correlation coefficient between the two trends. Hereafter, we will refer to it as SWFC– behavior correlation (SBC). Each seed ROI will thus get one SBC value. However, because of the very noisy characteristics of both fMRI and behavioral data, getting a significant SBC for every participant is problematic. Instead, we focused on the group-level SBC for which a subsample of 70% of the participants from each group were selected by simple random sampling without replacement. The group-averaged SWFC and illusory /ta/ perception trends were estimated for each subsample. The SBC for was then calculated as the Pearson correlation between these two averaged trends. Finally, the sampling was repeated 50 times, thus generating 50 SBC values for each seed ROI. The reported seed SBC is the averaged significant correlation value from the 50 subsamples (one-sample t-test; *p* ⩽ 0.05, FDR corrected). Taking 50 subsamples was necessary because of the experimental design where we had only 6 lag conditions (6 data points) on which the correlation was calculated, and the subsamples thus created the necessary distribution from which a population mean could be estimated.

#### Whole-brain graph theoretic network analysis

The required BOLD time series from 210 ROIs were extracted using REX toolbox (Whitfield-Gabrieli, 2009) (https://web.mit.edu/swg/software.htm). From each time series, task blocks were extracted based on the experimental design and then concatenated, and also further preprocessed through linear detrending (linear or linear breakpoint detrending). The functional interaction across all the 210 brain parcels were estimated using pairwise Pearson correlation between corresponding task-block BOLD time series (Lee Rodgers and Nicewander, 1988; Pearson, 1895). At the group level, the functional connectomes were generated as follows. First, distinct subsamples were generated by simple random sampling 70% of the total participants in each group. For each subsample, the averaged cross-correlation matrix was then evaluated incorporating Fisher’s z-transformation, and tested for statistical significance through one-sample t-test (*p* ⩽ 0.05, FDR corrected). We then applied a threshold of 0.48 to obtain the binarized network matrix; the threshold value of 0.48 was chosen to ensure a statistical power of ≈0.9. From those generated distinct subsamples, we finally selected 50 binarized networks which form a single connected component, i.e., there are no disconnected nodes in these networks. This condition was necessary to allow comparisons across the same network size, as the binarized networks generated with a fixed thresholding sometimes produced isolated nodes (or groups of isolated nodes). Out of the total 210 nodes, one node in the anterior right parahippocampal area that was frequently disconnected irrespective of stimulus conditions was excluded from the network study. Hence, the network size of the finally generated 50 brain networks was 209 × 209. Furthermore, for each generated binarized brain network we also generated 50 randomized connected network samples with same degree sequence. We used the MATLAB script *randmio und connected*.*m* available with Brain Connectivity Toolbox (BCT) for the randomization, keeping edge iterations to 30 (Rubinov and Sporns, 2010). The randomized networks were used to estimate the normalized network parameters, for a better comparison between networks of same sizes (especially when network size > 200 nodes) (Van Wijk et al., 2010).

To study the functional integration and segregation at different scales of brain network, we calculated the normalized average local clustering coefficient (*C*_n_), characteristics path length (*L*_n_), modularity (*Q*_n_), and node participation coefficients (*P*). *C*_n_ and *L*_n_ estimate the network characteristics at the resolution of nodes (local scale), and *Q*_n_ and *P* estimates at the level of the entire network (global scale). The estimation of modularity and participation coefficients required partitioning each brain network into nonoverlapping functional modules. For this purpose we used Blondel et al. (Blondel et al., 2008) method (Louvain method; *community louvain*.*m* code available with BCT). The graph measures were then normalized and compared across networks (Guimera et al., 2007; Van Wijk et al., 2010). See also Supplementary Methods.

## Acknowledgments

This study was supported by NBRC Core funds and Computing Facility. AB was supported by Ramalingaswami Fellowship (BT/RLF/Re-entry/31/2011) and Innovative Young Biotechnologist Award (IYBA, BT/07/IYBA/2013), both from Department of Biotechnology, Ministry of Science and Technology, Government of India. SSS’s and DR’s research was made possible through the financial support provided by Ashoka university and also by Axis Bank. SSS was also supported by a National Post-Doctoral Fellowship from Science and Engineering Research Board, Government of India (SERB-NPDF; No. PDF/2016/003188).

