## Supplementary Information for "High segregation and diminished global integration in large-scale brain functional networks enhances the perceptual binding of cross-modal stimuli"

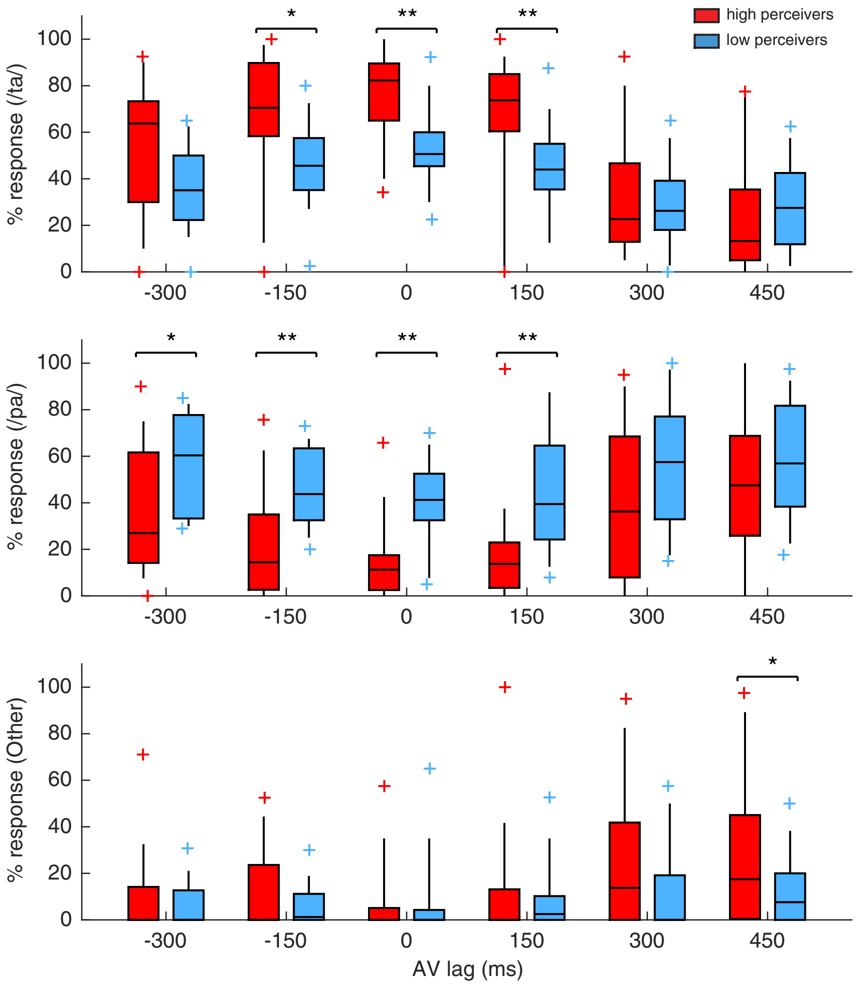


**Supplementary Figure 1.** Box plots showing the variation in /ta/, /pa/, and "Other" perceptual responses between high and low perceivers. Significant difference with *$p < 0.05$, **$p < 0.005$ (Welch's t-test).

A

B

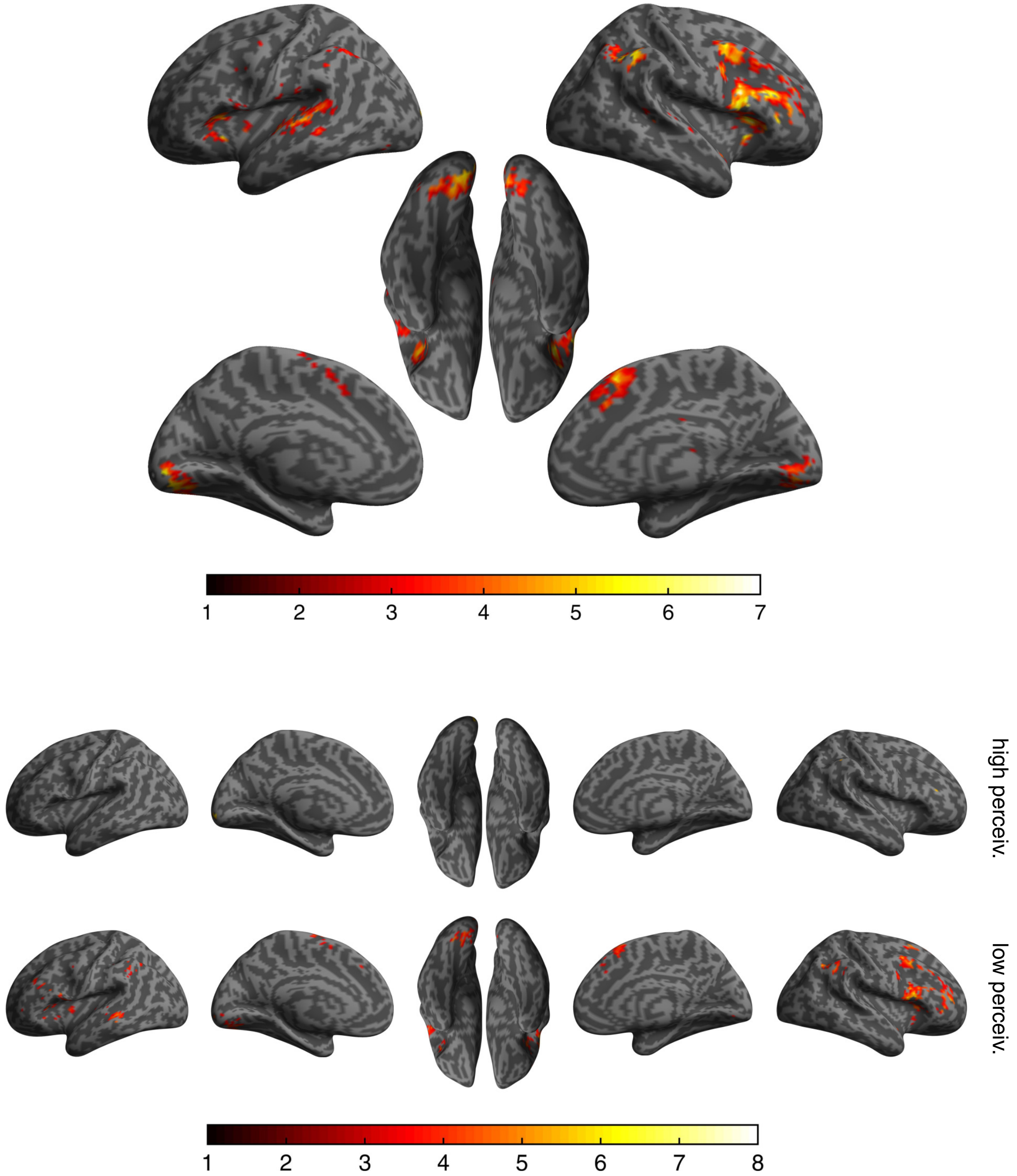


**Supplementary Figure 2.** Group-level task-enhanced (task $>$ rest) active brain areas (**A**) for all the 38 participants and (**B**) separately for high and low perceivers during Pure /ta/ stimulus condition (peak $p \leqslant 0.05$, FDR corrected). Color bar represents the T-score. Further, the active clusters from all the 38 participants taken together (panel A) are mapped on the Brainnetome atlas to identify the 29 seed ROIs used in this study (see also Supplementary Table 3).

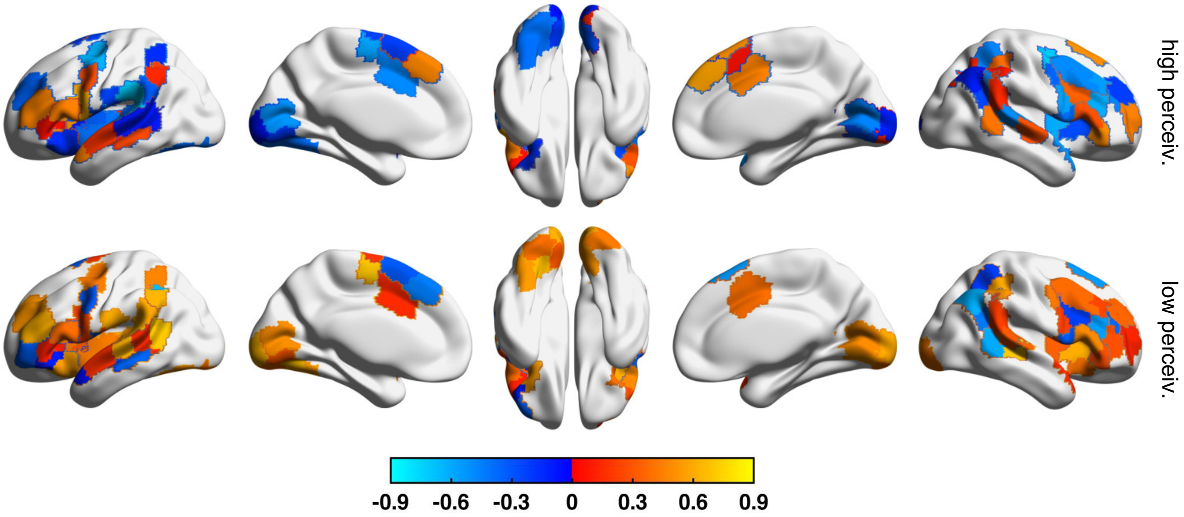


**Supplementary Figure 3.** Brain surface plots showing the SWFC–behavior correlation (SBC) changes of the seed regions between the high perceivers (top) and low perceivers (bottom), based on the parcellation scheme of the Schaefer atlas. Only the seeds with significant correlation are shown ($p < 0.05$, FDR corrected).

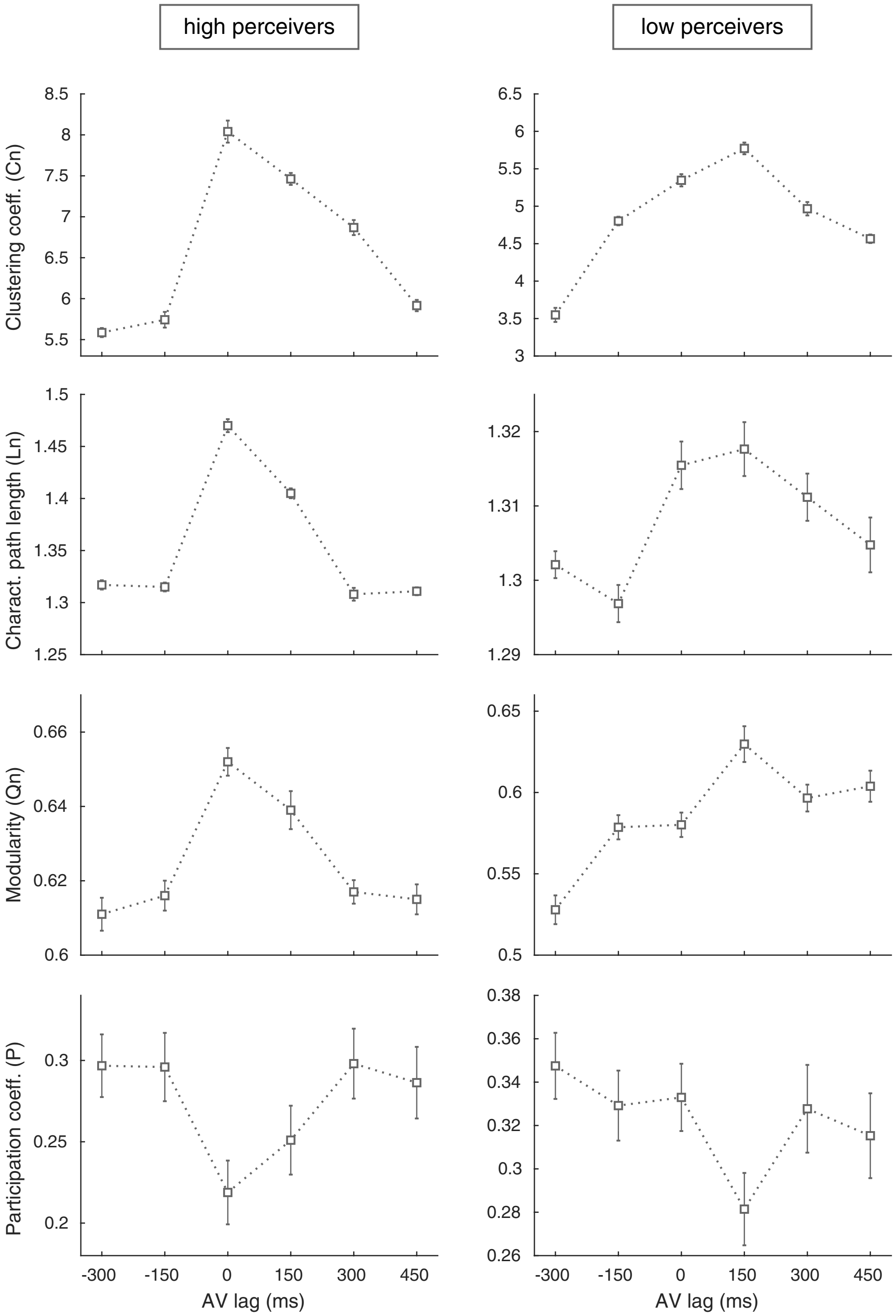


**Supplementary Figure 4.** Plot showing the changes in network topological measures for high (left column) and low perceivers (right column) across AV lag conditions, based on the parcellation scheme of the Schaefer atlas.

**Supplementary Table 1. Cluster with significant main effects of AV lags and grouping, and the their interaction effect ($p \leqslant 0.05$, FDR corrected).**

| **Left hemisphere** | | | | | **Right hemisphere** | | | | |
|---|---|---|---|---|---|---|---|---|---|
| **Brainnetome ROI label** | **Brain area** | **No. of voxels (cluster)** | ***F* score** | **Peak MNI coordinates** | **Brainnetome ROI label** | **Brain area** | **No. of voxels (cluster)** | ***F* score** | **Peak MNI coordinates** |
| ***AV Lag Effect*** | | | | | | | | | |
| SFG_L_7_1 | medial area 8 | 3 | 2.54 | [0,24,46] | SFG_R_7_1 | medial area 8 | 8 | 2.69 | [2,24,48] |
| | | 5 | 2.39 | [-8,16,54] | SFG_R_7_2 | dorsolateral area 8 | 2 | 2.48 | [18,28,42] |
| SFG_L_7_5 | medial area 6 | 5 | 2.43 | [-10,4,40] | SFG_R_7_6 | medial area 9 | 1 | 2.54 | [2,26,48] |
| SFG_L_7_6 | medial area 9 | 6 | 2.88 | [-6,26,34] | MFG_R_7_2 | inferior frontal junction | 2 | 2.29 | [48,22,30] |
| MFG_L_7_2 | inferior frontal junction | 6 | 2.62 | [-34,16,30] | MFG_R_7_4 | ventral area 9/46 | 3 | 2.47 | [36,44,16] |
| IFG_L_6_2 | inferior frontal sulcus | 72 | 3.42 | [-46,36,4] | MFG_R_7_5 | ventrolateral area 8 | 2 | 2.44 | [48,24,30] |
| IFG_L_6_3 | caudal area 45 | 3 | 2.53 | [-50,28,6] | IFG_R_6_1 | dorsal area 44 | 6 | 2.81 | [48,22,20] |
| IFG_L_6_4 | rostral area 45 | 22 | 4.02 | [-46,36,2] | | | 3 | 2.59 | [50,24,28] |
| IFG_L_6_5 | opercular area 44 | 3 | 2.75 | [-42,34,-2] | | | 2 | 2.58 | [46,14,20] |
| IFG_L_6_6 | ventral area 44 | 36 | 3.77 | [-54,10,2] | IFG_R_6_2 | inferior frontal sulcus | 40 | 4.03 | [46,34,10] |
| PrG_L_6_1 | area 4 (head/face) | 2 | 2.96 | [-60,8,18] | | | 7 | 3.16 | [46,24,18] |
| PrG_L_6_3 | area 4 (upper limb region) | 4 | 2.47 | [-26,-26,50] | IFG_R_6_3 | caudal area 45 | 3 | 3.8 | [48,16,8] |
| PrG_L_6_5 | area 4 (tongue/larynx) | 22 | 4.05 | [-54,8,0] | | | 2 | 2.34 | [50,20,16] |
| PrG_L_6_6 | caudal ventrolateral area 6 | 3 | 2.95 | [-54,8,4] | IFG_R_6_5 | opercular area 44 | 4 | 3.16 | [46,28,6] |
| | | 6 | 2.59 | [-58,10,14] | | | 2 | 2.48 | [46,16,6] |
| STG_L_6_2 | area 41/42 | 28 | 3.71 | [-42,-38,16] | IFG_R_6_6 | ventral area 44 | 1 | 2.64 | [46,14,18] |
| | | 10 | 2.44 | [-58,-40,20] | STG_R_6_2 | area 41/42 | 6 | 2.74 | [58,-28,16] |
| STG_L_6_3 | TE1.0 and TE1.2 | 15 | 3.06 | [-56,6,-2] | STG_R_6_3 | TE1.0 and TE1.2 | 14 | 2.75 | [48,-10,2] |
| | | 11 | 2.72 | [-42,-22,6] | | | 5 | 2.59 | [50,8,-10] |
| STG_L_6_4 | caudal area 22 | 22 | 3.83 | [-62,-44,14] | STG_R_6_5 | lateral area 38 | 1 | 2.29 | [42,-6,-12] |
| | | 11 | 3 | [-58,-24,-4] | SPL_R_5_2 | caudal area 7 | 15 | 3.19 | [26,-66,54] |
| | | 8 | 2.76 | [-66,-30,2] | SPL_R_5_3 | lateral area 5 | 5 | 2.45 | [38,-38,56] |
| STG_L_6_5 | lateral area 38 | 8 | 3.3 | [-46,12,-20] | SPL_R_5_5 | intraparietal area 7 (hIP3) | 2 | 2.43 | [26,-62,54] |
| STG_L_6_6 | rostral area 22 | 26 | 3.29 | [-60,-2,-4] | IPL_R_6_1 | caudal area 39 (PGp) | 6 | 2.89 | [34,-74,18] |
| | | 10 | 2.92 | [-50,-18,-6] | | | 2 | 2.64 | [40,-60,8] |
| MTG_L_4_3 | dorsolateral area 37 | 18 | 3.25 | [-56,-56,0] | IPL_R_6_6 | rostroventral area 40 (PFop) | 39 | 4.26 | [58,-30,22] |
| | | 8 | 2.75 | [-52,-68,2] | PCun_R_4_3 | parietooccipital sulcus (PEr) | 1 | 2.77 | [2,-58,10] |
| MTG_L_4_4 | anterior STS | 6 | 3.17 | [-54,-28,-4] | | | 1 | 2.51 | [24,-62,6] |
| | | 19 | 2.94 | [-66,-30,0] | INS_R_6_1 | hypergranular insula | 35 | 4.24 | [40,-16,2] |
| ITG_L_7_2 | lateroventral area 37 | 12 | 3.55 | [-48,-52,-14] | | | 4 | 4.04 | [42,-4,-4] |
| | | 3 | 3.08 | [-50,-66,-10] | INS_R_6_4 | v. dysgranular and granular | 37 | 4.61 | [40,-6,-6] |
| | | 3 | 2.42 | [-52,-60,-10] | | | 7 | 3.26 | [36,-16,-6] |
| ITG_L_7_5 | ventrolateral area 37 | 83 | 4.87 | [-52,-68,-6] | | | 3 | 2.71 | [44,8,-10] |
| FuG_L_3_1 | rostroventral area 20 | 3 | 2.72 | [-26,-40,-22] | INS_R_6_5 | dorsal granular insula | 14 | 4.12 | [38,-12,8] |
| FuG_L_3_2 | medioventral area37 | 5 | 2.75 | [-34,-58,-12] | | | 21 | 3.68 | [40,-4,-2] |
| | | 6 | 2.42 | [-32,-44,-20] | INS_R_6_6 | dorsal dysgranular insula | 3 | 2.48 | [38,4,0] |
| FuG_L_3_3 | lateroventral area37 | 176 | 4.5 | [-46,-54,-16] | MVOcC_R_5_2 | rostral cuneus gyrus | 4 | 2.91 | [2,-60,8] |
| pSTS_L | posterior STS | 29 | 3.23 | [-50,-40,8] | MVOcC_R_5_5 | vm. parietooccipital sulcus | 4 | 2.52 | [22,-62,6] |
| SPL_L_5_2 | caudal area 7 | 11 | 2.53 | [-24,-60,46] | | | 4 | 2.33 | [4,-58,6] |
| SPL_L_5_3 | lateral area 5 | 35 | 3.2 | [-36,-42,50] | | | 1 | 2.55 | [2,-60,10] |
| SPL_L_5_5 | intraparietal area 7 (hIP3) | 7 | 2.54 | [-26,-60,48] | LOcC_R_4_1 | middle occipital gyrus | 14 | 2.7 | [34,-82,16] |
| IPL_L_6_3 | rostrodorsal area 40 (PFt) | 66 | 3.36 | [-56,-28,40] | | | 1 | 3.17 | [36,-76,8] |
| | | 31 | 3.2 | [-40,-40,48] | LOcC_R_2_2 | lateral sup. occipital gyrus | 19 | 3.01 | [32,-74,16] |
| IPL_L_6_4 | caudal area 40 (PFm) | 34 | 3.75 | [-54,-46,28] | Tha_R | thalamus | 49 | 3.15 | [10,-20,6] |
| IPL_L_6_5 | rostroventral area 39 (PGa) | 5 | 2.53 | [-44,-48,20] | | | | | |
| | | 3 | 2.33 | [-48,-68,14] | | | | | |
| IPL_L_6_6 | rostroventral area 40 (PFop) | 268 | 4.29 | [-60,-34,30] | | | | | |
| PCun_L_4_3 | parietooccipital sulcus (PEr) | 3 | 2.49 | [-2,-60,8] | | | | | |
| PCun_L_4_4 | area 31 (Lc1) | 1 | 2.31 | [0,-58,12] | | | | | |
| PoG_L_4_3 | area 2 | 65 | 3.47 | [-38,-40,50] | | | | | |
| INS_L_6_1 | hypergranular insula | 4 | 3.15 | [-42,-10,-2] | | | | | |
| | | 2 | 2.42 | [-36,-16,0] | | | | | |
| INS_L_6_4 | v. dysgranular and granular | 69 | 3.09 | [-40,-10,-4] | | | | | |
| INS_L_6_5 | dorsal granular insula | 1 | 2.3 | [-38,0,-4] | | | | | |
| INS_L_6_6 | dorsal dysgranular insula | 2 | 2.74 | [-38,2,-4] | | | | | |
| CG_L_7_3 | pregenual area 32 | 1 | 2.26 | [-6,24,32] | | | | | |
| CG_L_7_5 | caudodorsal area 24 | 7 | 2.77 | [-10,6,34] | | | | | |
| CG_L_7_6 | caudal area 23 | 10 | 3.39 | [-10,-16,38] | | | | | |
| LOcC_L_4_2 | area V5/MT+ | 191 | 5.04 | [-50,-70,-8] | | | | | |
| LOcC_L_2_2 | lateral sup. occipital gyrus | 4 | 2.55 | [-24,-68,44] | | | | | |
| Tha_L | thalamus | 5 | 2.52 | [-8,-22,6] | | | | | |
| | | 7 | 2.45 | [-22,-22,4] | | | | | |

*Continued...*

| Left hemisphere | | | | | Right hemisphere | | | | |
|---|---|---|---|---|---|---|---|---|---|
| Brainnetome ROI label | Brain area | No. of voxels (cluster) | *F* score | Peak MNI coordinates | Brainnetome ROI label | Brain area | No. of voxels (cluster) | *F* score | Peak MNI coordinates |
| ***Group Effect*** | | | | | | | | | |
| SFG_L_7_1 | medial area 8 | 21 | 13.63 | [-6,24,52] | SFG_R_7_1 | medial area 8 | 58 | 17.12 | [8,24,56] |
| | | 4 | 10.51 | [-10,14,60] | SFG_R_7_2 | dorsolateral area 8 | 175 | 25.7 | [24,22,52] |
| SFG_L_7_2 | dorsolateral area 8 | 369 | 37.01 | [-24,26,44] | SFG_R_7_3 | lateral area 9 | 111 | 20.03 | [16,36,50] |
| SFG_L_7_3 | lateral area 9 | 132 | 39.31 | [-12,44,46] | | | 2 | 8.46 | [4,46,44] |
| | | 20 | 30.03 | [-16,58,22] | SFG_R_7_4 | dorsolateral area 6 | 37 | 20.08 | [24,-2,48] |
| SFG_L_7_4 | dorsolateral area 6 | 61 | 18.4 | [-24,8,62] | SFG_R_7_5 | medial area 6 | 25 | 10.08 | [4,-10,56] |
| SFG_L_7_5 | medial area 6 | 3 | 8.4 | [0,-12,56] | SFG_R_7_6 | medial area 9 | 50 | 15.92 | [2,38,48] |
| SFG_L_7_6 | medial area 9 | 228 | 26.98 | [-8,38,28] | | | 41 | 15.57 | [0,44,26] |
| SFG_L_7_7 | medial area 10 | 185 | 26.43 | [-8,38,26] | | | 8 | 9.74 | [8,38,34] |
| MFG_L_7_1 | dorsal area 9/46 | 52 | 26.5 | [-32,36,28] | | | 6 | 8.84 | [2,22,40] |
| | | 52 | 25.99 | [-20,54,24] | | | 2 | 8.83 | [0,32,38] |
| | | 42 | 18.05 | [-24,38,44] | SFG_R_7_7 | medial area 10 | 12 | 18.11 | [8,54,12] |
| MFG_L_7_2 | inferior frontal junction | 16 | 9.89 | [-46,24,28] | | | 32 | 11.98 | [0,54,20] |
| MFG_L_7_3 | area 46 | 59 | 31.41 | [-18,58,22] | MFG_R_7_1 | dorsal area 9/46 | 146 | 22.18 | [34,32,34] |
| | | 13 | 25.96 | [-32,46,12] | MFG_R_7_2 | inferior frontal junction | 19 | 17.88 | [40,14,46] |
| MFG_L_7_4 | ventral area 9/46 | 25 | 24.5 | [-32,44,10] | | | 14 | 17.21 | [42,20,30] |
| | | 2 | 9.3 | [-46,26,30] | | | 8 | 9.87 | [48,16,40] |
| | | 10 | 9.27 | [-34,34,20] | MFG_R_7_3 | area 46 | 6 | 9.39 | [24,46,24] |
| MFG_L_7_5 | ventrolateral area 8 | 223 | 36.11 | [-24,26,42] | MFG_R_7_4 | ventral area 9/46 | 43 | 31.3 | [38,24,24] |
| | | 6 | 13.22 | [-34,14,52] | | | 19 | 12.55 | [40,46,16] |
| MFG_L_7_6 | ventrolateral area 6 | 78 | 22.66 | [-26,8,60] | | | 5 | 9.19 | [42,34,18] |
| IFG_L_6_1 | dorsal area 44 | 5 | 11.91 | [-40,10,20] | MFG_R_7_5 | ventrolateral area 8 | 8 | 20.16 | [38,26,26] |
| IFG_L_6_3 | caudal area 45 | 10 | 14.9 | [-50,22,0] | | | 5 | 12.16 | [36,30,34] |
| IFG_L_6_4 | rostral area 45 | 2 | 18.61 | [-48,24,-4] | | | 2 | 11.12 | [40,16,46] |
| | | 2 | 10.41 | [-50,30,2] | MFG_R_7_6 | ventrolateral area 6 | 66 | 19.41 | [26,-2,48] |
| IFG_L_6_5 | opercular area 44 | 66 | 31.82 | [-32,26,8] | IFG_R_6_1 | dorsal area 44 | 73 | 32.57 | [38,22,22] |
| | | 45 | 19.7 | [-48,24,-2] | IFG_R_6_2 | inferior frontal sulcus | 2 | 23.39 | [40,24,20] |
| IFG_L_6_6 | ventral area 44 | 31 | 16.19 | [-50,12,10] | | | 2 | 8.26 | [44,34,20] |
| | | 6 | 15.58 | [-50,20,-2] | IFG_R_6_3 | caudal area 45 | 1 | 7.86 | [46,20,12] |
| OrG_L_6_2 | orbital area 12/47 | 4 | 10.16 | [-40,18,-14] | IFG_R_6_5 | opercular area 44 | 44 | 17.33 | [32,28,6] |
| | | 2 | 8.83 | [-34,36,-6] | | | 4 | 11.39 | [38,16,12] |
| OrG_L_6_6 | lateral area 12/47 | 24 | 20.67 | [-30,26,4] | IFG_R_6_6 | ventral area 44 | 13 | 12.26 | [50,12,18] |
| | | 7 | 14.61 | [-46,24,-6] | OrG_R_6_6 | lateral area 12/47 | 9 | 13.33 | [32,28,2] |
| | | 5 | 9.38 | [-40,20,-12] | PrG_R_6_1 | area 4 (head/face) | 3 | 9.47 | [46,-6,34] |
| PrG_L_6_1 | area 4 (head/face) | 147 | 22.22 | [-34,-18,40] | PrG_R_6_2 | caudal dorsolateral area 6 | 6 | 18.77 | [26,-4,48] |
| PrG_L_6_2 | caudal dorsolateral area 6 | 14 | 9.59 | [-28,-12,52] | PrG_R_6_5 | area 4 (tongue/larynx) | 3 | 16.21 | [52,8,0] |
| PrG_L_6_3 | area 4 (upper limb region) | 83 | 14.51 | [-24,-30,52] | | | 4 | 9.3 | [54,-2,6] |
| PrG_L_6_4 | area 4 (trunk region) | 5 | 16.46 | [-18,-22,68] | PrG_R_6_6 | caudal ventrolateral area 6 | 2 | 9.64 | [54,12,18] |
| PrG_L_6_5 | area 4 (tongue/larynx) | 29 | 22.57 | [-52,-6,6] | PCL_R_2_1 | area 1/2/3 (lower limb region) | 18 | 17.7 | [16,-42,52] |
| | | 2 | 7.97 | [-46,0,0] | PCL_R_2_2 | area 4 (lower limb region) | 37 | 10.68 | [2,-12,56] |
| PrG_L_6_6 | caudal ventrolateral area 6 | 8 | 12.39 | [-46,0,22] | | | 2 | 7.57 | [6,-12,66] |
| | | 5 | 12.18 | [-58,8,32] | STG_R_6_2 | area 41/42 | 25 | 15.51 | [50,-16,4] |
| | | 7 | 9.31 | [-44,2,40] | STG_R_6_3 | TE1.0 and TE1.2 | 30 | 16.48 | [52,-14,4] |
| PCL_L_2_1 | area1/2/3 (lower limb region) | 7 | 23.3 | [-18,-38,42] | | | 10 | 13.7 | [52,-6,6] |
| PCL_L_2_2 | area 4 (lower limb region) | 18 | 10.58 | [-8,-20,50] | | | 10 | 11.68 | [52,6,-2] |
| | | 3 | 8.11 | [-2,-32,60] | | | 2 | 8.57 | [44,-4,-12] |
| STG_L_6_1 | medial area 38 | 5 | 13.99 | [-44,14,-32] | STG_R_6_5 | lateral area 38 | 38 | 14.44 | [44,16,-16] |
| STG_L_6_2 | area 41/42 | 60 | 33.59 | [-56,-22,4] | | | 2 | 8.37 | [42,-4,-14] |
| | | 5 | 13.09 | [-58,-34,10] | FuG_R_3_1 | rostroventral area 20 | 48 | 39.18 | [30,-40,-14] |
| | | 12 | 10.61 | [-40,-40,14] | FuG_R_3_2 | medioventral area 37 | 108 | 37.36 | [30,-42,-14] |
| STG_L_6_3 | TE1.0 and TE1.2 | 156 | 31.6 | [-54,-22,6] | PhG_R_6_3 | l. parahippocampal gyrus | 46 | 25.24 | [32,-36,-12] |
| | | 2 | 11.71 | [-46,10,-10] | PhG_R_6_6 | m. parahippocampal gyrus | 61 | 24.17 | [28,-38,-12] |
| | | 5 | 9.87 | [-44,-22,0] | pSTS_R | posterior STS | 7 | 9.5 | [44,-42,14] |
| STG_L_6_4 | caudal area 22 | 43 | 18.1 | [-52,-24,0] | SPL_R_5_1 | rostral area 7 | 11 | 10.83 | [12,-58,62] |
| | | 12 | 10.63 | [-62,-32,6] | | | 3 | 8.27 | [22,-54,56] |
| | | 2 | 7.57 | [-60,-44,16] | SPL_R_5_3 | lateral area 5 | 4 | 9.94 | [28,-38,54] |
| STG_L_6_5 | lateral area 38 | 21 | 26.99 | [-46,16,-26] | SPL_R_5_4 | postcentral area 7 | 3 | 10.07 | [26,-38,54] |
| | | 19 | 15.41 | [-42,14,-14] | IPL_R_6_5 | rostroventral area 39 (PGa) | 4 | 9.46 | [40,-54,22] |
| MTG_L_4_2 | rostral area 21 | 3 | 16.12 | [-44,12,-32] | IPL_R_6_6 | rostroventral area 40 (PFop) | 46 | 12.98 | [48,-20,20] |
| MTG_L_4_3 | dorsolateral area 37 | 5 | 15.15 | [-56,-52,2] | PCun_R_4_1 | medial area 7 (PEp) | 62 | 17.69 | [6,-64,50] |
| | | 8 | 9.01 | [-56,-60,0] | PCun_R_4_2 | medial area 5 (PEm) | 166 | 24.07 | [14,-44,52] |
| MTG_L_4_4 | anterior STS | 1 | 7.84 | [-54,-28,-4] | PCun_R_4_3 | parietooccipital sulcus (PEr) | 226 | 36.41 | [20,-72,18] |
| ITG_L_7_2 | lateroventral area 37 | 13 | 13.17 | [-50,-52,-14] | | | 10 | 13.05 | [18,-68,26] |
| ITG_L_7_5 | ventrolateral area 37 | 65 | 17.1 | [-52,-56,-6] | | | 9 | 11.64 | [20,-78,28] |
| ITG_L_7_6 | caudolateral of area 20 | 1 | 9.4 | [-54,-52,-10] | PCun_R_4_4 | area 31 (Lc1) | 8 | 16.86 | [6,-54,46] |
| FuG_L_3_1 | rostroventral area 20 | 98 | 36.66 | [-26,-46,-12] | | | 11 | 11.61 | [6,-52,14] |

*Continued...*

| Left hemisphere | | | | |
|---|---|---|---|---|
| Brainnetome ROI label | Brain area | No. of voxels (cluster) | *F* score | Peak MNI coordinates |
| FuG_L_3_2 | medioventral area 37 | 221 | 42.33 | [-26,-46,-14] |
| | | 27 | 21.29 | [-22,-76,-16] |
| | | 68 | 17.76 | [-38,-70,-18] |
| | | 5 | 10.66 | [-38,-54,-20] |
| FuG_L_3_3 | lateroventral area37 | 19 | 17.7 | [-40,-54,-18] |
| | | 4 | 12.31 | [-40,-70,-16] |
| | | 2 | 8.95 | [-48,-56,-16] |
| PhG_L_6_3 | l. parahippocampal gyrus | 44 | 26.49 | [-28,-40,-14] |
| PhG_L_6_6 | m. parahippocampal gyrus | 58 | 34.63 | [-20,-44,-14] |
| pSTS_L | posterior STS | 36 | 25.17 | [-54,-34,6] |
| | | 153 | 22.74 | [-54,-50,4] |
| SPL_L_5_1 | rostral area 7 | 10 | 10.62 | [-10,-62,62] |
| | | 15 | 9.58 | [-20,-54,56] |
| SPL_L_5_2 | caudal area 7 | 33 | 10.43 | [-22,-68,48] |
| SPL_L_5_3 | lateral area 5 | 23 | 13.45 | [-28,-40,46] |
| | | 16 | 9.92 | [-32,-44,56] |
| | | 2 | 7.97 | [-32,-46,36] |
| SPL_L_5_4 | postcentral area 7 | 38 | 12.38 | [-18,-42,68] |
| | | 8 | 8.95 | [-30,-44,58] |
| SPL_L_5_5 | intraparietal area 7 (hIP3) | 23 | 11.04 | [-24,-60,54] |
| IPL_L_6_1 | caudal area 39 (PGp) | 248 | 52.58 | [-40,-82,22] |
| | | 48 | 14.82 | [-26,-70,38] |
| IPL_L_6_2 | rostrodorsal area 39 (Hip3) | 59 | 23.31 | [-40,-66,36] |
| | | 2 | 8.35 | [-44,-50,40] |
| IPL_L_6_3 | rostrodorsal area 40 (PFt) | 26 | 12.56 | [-36,-32,38] |
| IPL_L_6_4 | caudal area 40 (PFm) | 120 | 19.86 | [-48,-58,36] |
| | | 7 | 10.31 | [-56,-44,30] |
| IPL_L_6_5 | rostroventral area 39 (PGa) | 463 | 36.96 | [-44,-76,32] |
| IPL_L_6_6 | rostroventral area 40 (PFop) | 19 | 12.07 | [-42,-40,24] |
| | | 4 | 11.62 | [-54,-22,12] |
| | | 4 | 10.85 | [-56,-44,28] |
| | | 6 | 10.34 | [-42,-24,22] |
| PCun_L_4_1 | medial area 7 (PEp) | 7 | 12.02 | [-10,-64,48] |
| | | 4 | 11.29 | [-8,-58,44] |
| | | 3 | 8.34 | [0,-64,46] |
| | | 3 | 7.79 | [-10,-60,58] |
| PCun_L_4_2 | medial area 5 (PEm) | 48 | 26.45 | [-16,-46,50] |
| | | 11 | 24.18 | [-16,-40,44] |
| | | 3 | 9 | [-2,-48,54] |
| PCun_L_4_3 | parietooccipital sulcus (PEr) | 234 | 32.08 | [-10,-66,18] |
| | | 25 | 12.71 | [-10,-74,36] |
| PCun_L_4_4 | area 31 (Lc1) | 10 | 22.72 | [-8,-54,44] |
| | | 27 | 18.81 | [-12,-54,34] |
| | | 10 | 12.02 | [-6,-54,16] |
| PoG_L_4_1 | area 1/2/3 (limb/head/face) | 75 | 17.91 | [-58,-10,32] |
| | | 53 | 16.88 | [-38,-18,44] |
| PoG_L_4_2 | area 1/2/3 (tongue/larynx) | 14 | 14.64 | [-54,-10,8] |
| PoG_L_4_3 | area 2 | 13 | 12.65 | [-36,-32,42] |
| | | 4 | 11.3 | [-30,-38,62] |
| PoG_L_4_4 | area1/2/3 (trunk region) | 65 | 13.39 | [-26,-30,54] |
| INS_L_6_1 | hypergranular insula | 4 | 11.28 | [-40,-16,12] |
| INS_L_6_2 | ventral agranular insula | 5 | 9.94 | [-40,18,-12] |
| INS_L_6_3 | dorsal agranular insula | 63 | 28.16 | [-34,24,8] |
| | | 6 | 10.92 | [-42,14,-10] |
| INS_L_6_4 | v. dysgranular and granular | 20 | 12.98 | [-36,-6,-10] |
| INS_L_6_5 | dorsal granular insula | 7 | 10.71 | [-40,-14,12] |
| | | 7 | 10.34 | [-36,-4,-2] |
| INS_L_6_6 | dorsal dysgranular insula | 4 | 13.96 | [-44,12,-8] |
| | | 2 | 7.98 | [-44,2,0] |
| CG_L_7_1 | dorsal area 23 | 6 | 16.94 | [0,-26,38] |
| | | 14 | 10.99 | [0,-44,22] |
| CG_L_7_3 | pregenual area 32 | 50 | 23.81 | [-8,40,24] |
| CG_L_7_4 | ventral area 23 | 115 | 22.12 | [-14,-54,10] |
| CG_L_7_6 | caudal area 23 | 5 | 20.12 | [-16,-36,42] |
| | | 9 | 14.18 | [0,-24,38] |
| | | 10 | 9.8 | [-12,-18,36] |

| Right hemisphere | | | | |
|---|---|---|---|---|
| Brainnetome ROI label | Brain area | No. of voxels (cluster) | *F* score | Peak MNI coordinates |
| PoG_R_4_2 | area 1/2/3 (tongue/larynx) | 50 | 17.61 | [54,-14,14] |
| | | 14 | 14.07 | [50,-6,6] |
| PoG_R_4_3 | area 2 | 9 | 10.73 | [28,-36,54] |
| PoG_R_4_4 | area 1/2/3 (trunk region) | 13 | 9.61 | [24,-30,58] |
| INS_R_6_1 | hypergranular insula | 30 | 11.13 | [34,-22,18] |
| INS_R_6_3 | dorsal agranular insula | 21 | 15.06 | [30,22,10] |
| INS_R_6_4 | v. dysgranular and granular | 6 | 8.83 | [42,-4,-12] |
| INS_R_6_5 | dorsal granular insula | 2 | 8.24 | [38,-2,0] |
| INS_R_6_6 | dorsal dysgranular insula | 1 | 8.12 | [38,0,2] |
| CG_R_7_1 | dorsal area 23 | 12 | 14.01 | [2,-24,36] |
| | | 5 | 9.32 | [2,-44,22] |
| CG_R_7_2 | rostroventral area 24 | 3 | 14.54 | [4,-10,32] |
| CG_R_7_3 | pregenual area 32 | 11 | 14.16 | [2,40,20] |
| CG_R_7_4 | ventral area 23 | 82 | 29.13 | [8,-50,8] |
| | | 2 | 10.73 | [20,-42,-4] |
| CG_R_7_5 | caudodorsal area 24 | 7 | 9.11 | [4,6,34] |
| CG_R_7_6 | caudal area 23 | 14 | 21.82 | [12,-24,36] |
| | | 5 | 16.26 | [4,-22,36] |
| | | 4 | 9.66 | [2,-10,32] |
| CG_R_7_7 | subgenual area 32 | 24 | 18.32 | [6,38,14] |
| MVOcC_R_5_1 | caudal lingual gyrus | 31 | 23.26 | [6,-86,-2] |
| MVOcC_R_5_2 | rostral cuneus gyrus | 76 | 44.19 | [2,-80,20] |
| | | 15 | 12.44 | [2,-78,0] |
| | | 42 | 12.34 | [10,-70,2] |
| | | 2 | 12.07 | [16,-76,16] |
| MVOcC_R_5_3 | caudal cuneus gyrus | 54 | 32.34 | [6,-90,2] |
| | | 21 | 30.78 | [4,-82,20] |
| | | 10 | 10.89 | [10,-88,22] |
| MVOcC_R_5_4 | rostral lingual gyrus | 372 | 30.18 | [22,-46,-10] |
| MVOcC_R_5_5 | vm. parietooccipital sulcus | 179 | 34.63 | [16,-72,22] |
| | | 192 | 21.5 | [10,-50,4] |
| LOcC_R_4_1 | middle occipital gyrus | 2 | 8.83 | [26,-86,6] |
| LOcC_R_4_3 | occipital polar cortex | 3 | 11.51 | [16,-90,20] |
| LOcC_R_4_4 | inferior occipital gyrus | 8 | 12.82 | [22,-80,-8] |
| LOcC_R_2_1 | medial sup. occipital gyrus | 7 | 11.83 | [14,-90,20] |
| | | 4 | 8.35 | [18,-86,22] |
| Tha_R | thalamus | 87 | 15.57 | [8,-12,14] |

*Continued...*

| Left hemisphere | | | | | Right hemisphere | | | | |
|---|---|---|---|---|---|---|---|---|---|
| **Brainnetome ROI label** | **Brain area** | **No. of voxels (cluster)** | ***F* score** | **Peak MNI coordinates** | **Brainnetome ROI label** | **Brain area** | **No. of voxels (cluster)** | ***F* score** | **Peak MNI coordinates** |
| MVOcC_L_5_1 | caudal lingual gyrus | 159 | 46.87 | [-6,-86,-6] | | | | | |
| MVOcC_L_5_2 | rostral cuneus gyrus | 202 | 47.31 | [0,-80,22] | | | | | |
| | | 76 | 31.39 | [-6,-86,-2] | | | | | |
| MVOcC_L_5_3 | caudal cuneus gyrus | 78 | 42.03 | [-6,-88,-4] | | | | | |
| MVOcC_L_5_4 | rostral lingual gyrus | 375 | 35.69 | [-26,-46,-6] | | | | | |
| | | 20 | 20.88 | [-20,-74,-12] | | | | | |
| MVOcC_L_5_5 | vm. parietooccipital sulcus | 573 | 53.82 | [-12,-80,22] | | | | | |
| | | 4 | 10.49 | [-8,-78,36] | | | | | |
| LOcC_L_4_1 | middle occipital gyrus | 12 | 13 | [-40,-78,14] | | | | | |
| LOcC_L_4_2 | area V5/MT+ | 8 | 9.54 | [-42,-76,14] | | | | | |
| LOcC_L_4_4 | inferior occipital gyrus | 15 | 15.76 | [-36,-88,-12] | | | | | |
| LOcC_L_2_1 | medial sup. occipital gyrus | 44 | 36.05 | [-6,-86,26] | | | | | |
| LOcC_L_2_2 | lateral sup. occipital gyrus | 7 | 13.75 | [-24,-82,38] | | | | | |
| | | 34 | 10.84 | [-22,-70,46] | | | | | |
| Tha_L | thalamus | 63 | 24.34 | [-8,-10,14] | | | | | |
| ***Interaction Effect*** | | | | | | | | | |
| SFG_L_7_1 | medial area 8 | 13 | 2.6 | [-10,18,52] | SFG_R_7_1 | medial area 8 | 8 | 2.45 | [8,24,48] |
| SFG_L_7_2 | dorsolateral area 8 | 9 | 2.53 | [-12,20,56] | SFG_R_7_2 | dorsolateral area 8 | 11 | 2.96 | [22,24,54] |
| SFG_L_7_6 | medial area 9 | 8 | 3.51 | [-10,36,32] | | | 15 | 2.66 | [16,32,48] |
| MFG_L_7_4 | ventral area 9/46 | 1 | 2.27 | [-36,34,10] | SFG_R_7_3 | lateral area 9 | 2 | 2.31 | [16,36,48] |
| IFG_L_6_5 | opercular area 44 | 5 | 2.83 | [-36,34,2] | MFG_R_7_1 | dorsal area 9/46 | 5 | 2.78 | [24,30,32] |
| PrG_L_6_3 | area 4 (upper limb region) | 44 | 3.66 | [-24,-26,52] | IFG_R_6_3 | caudal area 45 | 1 | 2.36 | [44,24,12] |
| PCL_L_2_1 | area 1/2/3 (lower limb region) | 2 | 2.93 | [-18,-32,40] | IFG_R_6_5 | opercular area 44 | 1 | 2.62 | [42,24,12] |
| STG_L_6_2 | area 41/42 | 5 | 2.64 | [-40,-40,18] | PCL_R_2_2 | area 4 (lower limb region) | 9 | 2.46 | [8,-22,58] |
| ITG_L_7_2 | lateroventral area 37 | 8 | 3.59 | [-48,-50,-14] | PoG_R_4_4 | area 1/2/3 (trunk region) | 2 | 2.35 | [20,-30,62] |
| FuG_L_3_3 | lateroventral area37 | 11 | 3.38 | [-46,-52,-14] | INS_R_6_1 | hypergranular insula | 9 | 3.11 | [38,-12,6] |
| SPL_L_5_4 | postcentral area 7 | 1 | 2.29 | [-22,-38,56] | INS_R_6_5 | dorsal granular insula | 9 | 3.3 | [38,-12,8] |
| IPL_L_6_6 | rostroventral area 40 (PFop) | 60 | 3.28 | [-40,-34,20] | CG_R_7_4 | ventral area 23 | 2 | 2.45 | [8,-26,28] |
| PoG_L_4_4 | area 1/2/3 (trunk region) | 2 | 2.49 | [-26,-28,56] | Tha_R | thalamus | 14 | 3.19 | [20,-20,16] |
| | | 9 | 2.46 | [-16,-38,66] | | | 2 | 2.38 | [8,-22,16] |
| INS_L_6_1 | hypergranular insula | 2 | 2.75 | [-32,-34,18] | | | | | |
| CG_L_7_6 | caudal area 23 | 10 | 2.73 | [-16,-28,40] | | | | | |
| Tha_L | thalamus | 10 | 3.32 | [-16,-30,14] | | | | | |

**Supplementary Table 2. Details of 210 ROIs in our study, with reference to the Brainnetome atlas.**

| Regions | Gyrus/Sulcus | Subregion Anatomical Description | Left/Right Hemisphere | ROI Label* | MNI Coordinates |
|---|---|---|---|---|---|
| Frontal lobe | Superior frontal gyrus (SFG) | A8m, medial area 8 | left | SFG_L_7_1 | [-5 ,15,54] |
| | | | right | SFG_R_7_1 | [7,16,54] |
| | | A8dl, dorsolateral area 8 | left | SFG_L_7_2 | [-18,24,53] |
| | | | right | SFG_R_7_2 | [22,26,51] |
| | | A9l, lateral area 9 | left | SFG_L_7_3 | [-11,49,40] |
| | | | right | SFG_R_7_3 | [13,48,40] |
| | | A6dl, dorsolateral area 6 | left | SFG_L_7_4 | [-18,-1,65] |
| | | | right | SFG_R_7_4 | [20,4,64] |
| | | A6m, medial area 6 | left | SFG_L_7_5 | [-6,-5,58] |
| | | | right | SFG_R_7_5 | [7,-4,60] |
| | | A9m,medial area 9 | left | SFG_L_7_6 | [-5,36,38] |
| | | | right | SFG_R_7_6 | [6,38,35] |
| | | A10m, medial area 10 | left | SFG_L_7_7 | [-8,56,15] |
| | | | right | SFG_R_7_7 | [8,58,13] |
| | Middle frontal gyrus (MFG) | A9/46d, dorsal area 9/46 | left | MFG_L_7_1 | [-27,43,31] |
| | | | right | MFG_R_7_1 | [30,37,36] |
| | | IFJ, inferior frontal junction | left | MFG_L_7_2 | [-42,13,36] |
| | | | right | MFG_R_7_2 | [42,11,39] |
| | | A46, area 46 | left | MFG_L_7_3 | [-28,56,12] |
| | | | right | MFG_R_7_3 | [28,55,17] |
| | | A9/46v, ventral area 9/46 | left | MFG_L_7_4 | [-41,41,16] |
| | | | right | MFG_R_7_4 | [42,44,14] |
| | | A8vl, ventrolateral area 8 | left | MFG_L_7_5 | [-33,23,45] |
| | | | right | MFG_R_7_5 | [42,27,39] |
| | | A6vl, ventrolateral area 6 | left | MFG_L_7_6 | [-32,4,55] |
| | | | right | MFG_R_7_6 | [34,8,54] |
| | | A10l, lateral area10 | left | MFG_L_7_7 | [-26,60,-6] |
| | | | right | MFG_R_7_7 | [25,61,-4] |
| | Inferior frontal gyrus (IFG) | A44d,dorsal area 44 | left | IFG_L_6_1 | [-46,13,24] |
| | | | right | IFG_R_6_1 | [45,16,25] |
| | | IFS, inferior frontal sulcus | left | IFG_L_6_2 | [-47,32,14] |
| | | | right | IFG_R_6_2 | [48,35,13] |
| | | A45c, caudal area 45 | left | IFG_L_6_3 | [-53,23,11] |
| | | | right | IFG_R_6_3 | [54,24,12] |
| | | A45r, rostral area 45 | left | IFG_L_6_4 | [-49,36,-3] |
| | | | right | IFG_R_6_4 | [51,36,-1] |
| | | A44op, opercular area 44 | left | IFG_L_6_5 | [-39,23,4] |
| | | | right | IFG_R_6_5 | [42,22,3] |
| | | A44v, ventral area 44 | left | IFG_L_6_6 | [-52,13,6] |
| | | | right | IFG_R_6_6 | [54,14,11] |
| | Orbital gyrus (OrG) | A14m, medial area 14 | left | OrG_L_6_1 | [-7,54,-7] |
| | | | right | OrG_R_6_1 | [6,47,-7] |
| | | A12/47o, orbital area 12/47 | left | OrG_L_6_2 | [-36,33,-16] |
| | | | right | OrG_R_6_2 | [40,39,-14] |
| | | A11l, lateral area 11 | left | OrG_L_6_3 | [-23,38,-18] |
| | | | right | OrG_R_6_3 | [23,36,-18] |
| | | A11m, medial area 11 | left | OrG_L_6_4 | [-6,52,-19] |
| | | | right | OrG_R_6_4 | [6,57,-16] |
| | | A13, area 13 | left | OrG_L_6_5 | [-10,18,-19] |
| | | | right | OrG_R_6_5 | [9,20,-19] |
| | | A12/47l, lateral area 12/47 | left | OrG_L_6_6 | [-41,32,-9] |
| | | | right | OrG_R_6_6 | [42,31,-9] |

*Continued...*

| Regions | Gyrus/Sulcus | Subregion Anatomical Description | Left/Right Hemisphere | ROI Label* | MNI Coordinates |
|---|---|---|---|---|---|
| | Precentral gyrus (PrG) | A4hf, area 4 (head and face region) | left | PrG_L_6_1 | [-49,-8,39] |
| | | | right | PrG_R_6_1 | [55,-2,33] |
| | | A6cdl, caudal dorsolateral area 6 | left | PrG_L_6_2 | [-32,-9,58] |
| | | | right | PrG_R_6_2 | [33,-7,57] |
| | | A4ul, area 4 (upper limb region) | left | PrG_L_6_3 | [-26,-25,63] |
| | | | right | PrG_R_6_3 | [34,-19,59] |
| | | A4t, area 4 (trunk region) | left | PrG_L_6_4 | [-13,-20,73] |
| | | | right | PrG_R_6_4 | [15,-22,71] |
| | | A4tl, area 4 (tongue and larynx region) | left | PrG_L_6_5 | [-52,0,8] |
| | | | right | PrG_R_6_5 | [54,4,9] |
| | | A6cvl, caudal ventrolateral area 6 | left | PrG_L_6_6 | [-49,5,30] |
| | | | right | PrG_R_6_6 | [51,7,30] |
| | Paracentral lobule (PCL) | A1/2/3ll, area1/2/3 (lower limb region) | left | PCL_L_2_1 | [-8,-38,58] |
| | | | right | PCL_R_2_1 | [10,-34,54] |
| | | A4ll, area 4 (lower limb region) | left | PCL_L_2_2 | [-4,-23,61] |
| | | | right | PCL_R_2_2 | [5,-21,61] |
| Temporal lobe | Superior temporal gyrus (STG) | A38m, medial area 38 | left | STG_L_6_1 | [-32,14,-34] |
| | | | right | STG_R_6_1 | [31,15,-34] |
| | | A41/42, area 41/42 | left | STG_L_6_2 | [-54,-32,12] |
| | | | right | STG_R_6_2 | [54,-24,11] |
| | | TE1.0 and TE1.2 | left | STG_L_6_3 | [-50,-11,1] |
| | | | right | STG_R_6_3 | [51,-4,-1] |
| | | A22c, caudal area 22 | left | STG_L_6_4 | [-62,-33,7] |
| | | | right | STG_R_6_4 | [66,-20,6] |
| | | A38l, lateral area 38 | left | STG_L_6_5 | [-45,11,-20] |
| | | | right | STG_R_6_5 | [47,12,-20] |
| | | A22r, rostral area 22 | left | STG_L_6_6 | [-55,-3,-10] |
| | | | right | STG_R_6_6 | [56,-12,-5] |
| | Middle temporal gyrus (MTG) | A21c, caudal area 21 | left | MTG_L_4_1 | [-65,-30,-12] |
| | | | right | MTG_R_4_1 | [65,-29,-13] |
| | | A21r, rostral area 21 | left | MTG_L_4_2 | [-53,2,-30] |
| | | | right | MTG_R_4_2 | [51,6,-32] |
| | | A37dl, dorsolateral area 37 | left | MTG_L_4_3 | [-59,-58,4] |
| | | | right | MTG_R_4_3 | [60,-53,3] |
| | | aSTS, anterior superior temporal sulcus | left | MTG_L_4_4 | [-58,-20,-9] |
| | | | right | MTG_R_4_4 | [58,-16,-10] |
| | Inferior temporal gyrus (ITG) | A20iv, intermediate ventral area 20 | left | ITG_L_7_1 | [-45,-26,-27] |
| | | | right | ITG_R_7_1 | [46,-14,-33] |
| | | A37elv, extreme lateroventral area 37 | left | ITG_L_7_2 | [-51,-57,-15] |
| | | | right | ITG_R_7_2 | [53,-52,-18] |
| | | A20r, rostral area 20 | left | ITG_L_7_3 | [-43,-2,-41] |
| | | | right | ITG_R_7_3 | [40, 0,-43] |
| | | A20il, intermediate lateral area 20 | left | ITG_L_7_4 | [-56,-16,-28] |
| | | | right | ITG_R_7_4 | [55,-11,-32] |
| | | A37vl, ventrolateral area 37 | left | ITG_L_7_5 | [-55,-60,-6] |
| | | | right | ITG_R_7_5 | [54,-57,-8] |
| | | A20cl, caudolateral of area 20 | left | ITG_L_7_6 | [-59,-42,-16] |
| | | | right | ITG_R_7_6 | [61,-40,-17] |
| | | A20cv, caudoventral of area 20 | left | ITG_L_7_7 | [-55,-31,-27] |
| | | | right | ITG_R_7_7 | [54,-31,-26] |
| | Fusiform gyrus (FuG) | A20rv, rostroventral area 20 | left | FuG_L_3_1 | [-33,-16,-32] |
| | | | right | FuG_R_3_1 | [33,-15,-34] |
| | | A37mv, medioventral area 37 | left | FuG_L_3_2 | [-31,-64,-14] |
| | | | right | FuG_R_3_2 | [31,-62,-14] |
| | | A37lv, lateroventral area 37 | left | FuG_L_3_3 | [-42,-51,-17] |
| | | | right | FuG_R_3_3 | [43,-49,-19] |

*Continued...*

| Regions | Gyrus/Sulcus | Subregion Anatomical Description | Left/Right Hemisphere | ROI Label* | MNI Coordinates |
|---|---|---|---|---|---|
| | Parahippocampal gyrus (PhG) | A35/36r, rostral area 35/36 | left<br>right | PhG_L_6_1<br>PhG_R_6_1 | [-27,-7,-34]<br>[28,-8,-33] |
| | | A35/36c, caudal area 35/36 | left<br>right | PhG_L_6_2<br>PhG_R_6_2 | [-25,-25,-26]<br>[26,-23,-27] |
| | | TL, area TL (lateral PPHC, posterior parahippocampal gyrus) | left<br>right | PhG_L_6_3<br>PhG_R_6_3 | [-28,-32,-18]<br>[30,-30,-18] |
| | | A28/34, area 28/34 (EC, entorhinal cortex) | left<br>right | PhG_L_6_4<br>PhG_R_6_4 | [-19,-12,-30]<br>[19,-10,-30] |
| | | TI, area TI (temporal agranular insular cortex) | left<br>right | PhG_L_6_5<br>PhG_R_6_5 | [-23,2,-32]<br>[22,1,-36] |
| | | TH, area TH (medial PPHC) | left<br>right | PhG_L_6_6<br>PhG_R_6_6 | [-17,-39,-10]<br>[19,-36,-11] |
| | Posterior superior temporal sulcus (pSTS) | Posterior superior temporal sulcus | left<br>right | pSTS_L<br>pSTS_R | [-53,-45,8]<br>[55,-38,8] |
| Parietal lobe | Superior parietal lobule (SPL) | A7r, rostral area 7 | left<br>right | SPL_L_5_1<br>SPL_R_5_1 | [-16,-60,63]<br>[19,-57,65] |
| | | A7c, caudal area 7 | left<br>right | SPL_L_5_2<br>SPL_R_5_2 | [-15,-71,52]<br>[19,-69,54] |
| | | A5l, lateral area 5 | left<br>right | SPL_L_5_3<br>SPL_R_5_3 | [-33,-47,50]<br>[35,-42,54] |
| | | A7pc, postcentral area 7 | left<br>right | SPL_L_5_4<br>SPL_R_5_4 | [-22,-47,65]<br>[23,-43,67] |
| | | A7ip, intraparietal area 7 (hIP3) | left<br>right | SPL_L_5_5<br>SPL_R_5_5 | [-27,-59,54]<br>[31,-54,53] |
| | Inferior parietal lobule (IPL) | A39c, caudal area 39 (PGp) | left<br>right | IPL_L_6_1<br>IPL_R_6_1 | [-34,-80,29]<br>[45,-71,20] |
| | | A39rd, rostrodorsal area 39 (Hip3) | left<br>right | IPL_L_6_2<br>IPL_R_6_2 | [-38,-61,46]<br>[39,-65,44] |
| | | A40rd, rostrodorsal area 40 (PFt) | left<br>right | IPL_L_6_3<br>IPL_R_6_3 | [-51,-33,42]<br>[47,-35,45] |
| | | A40c, caudal area 40 (PFm) | left<br>right | IPL_L_6_4<br>IPL_R_6_4 | [-56,-49,38]<br>[57,-44,38] |
| | | A39rv, rostroventral area 39 (PGa) | left<br>right | IPL_L_6_5<br>IPL_R_6_5 | [-47,-65,26]<br>[53,-54,25] |
| | | A40rv, rostroventral area 40 (PFop) | left<br>right | IPL_L_6_6<br>IPL_R_6_6 | [-53,-31,23]<br>[55,-26,26] |
| | Precuneus (Pcun) | A7m, medial area 7 (PEp) | left<br>right | PCun_L_4_1<br>PCun_R_4_1 | [-5,-63,51]<br>[6,-65,51] |
| | | A5m, medial area 5 (PEm) | left<br>right | PCun_L_4_2<br>PCun_R_4_2 | [-8,-47,57]<br>[7,-47,58] |
| | | dmPOS, dorsomedial parietooccipital sulcus (PEr) | left<br>right | PCun_L_4_3<br>PCun_R_4_3 | [-12,-67,25]<br>[16,-64,25] |
| | | A31, area 31 (Lc1) | left<br>right | PCun_L_4_4<br>PCun_R_4_4 | [-6,-55,34]<br>[6,-54,35] |
| | Postcentral gyrus (PoG) | A1/2/3ulhf, area 1/2/3 (upper limb, head and face region) | left<br>right | PoG_L_4_1<br>PoG_R_4_1 | [-50,-16,43]<br>[50,-14,44] |
| | | A1/2/3tonIa, area 1/2/3 (tongue and larynx region) | left<br>right | PoG_L_4_2<br>PoG_R_4_2 | [-56,-14,16]<br>[56,-10,15] |
| | | A2, area 2 | left<br>right | PoG_L_4_3<br>PoG_R_4_3 | [-46,-30,50]<br>[48,-24,48] |
| | | A1/2/3tru, area1/2/3 (trunk region) | left<br>right | PoG_L_4_4<br>PoG_R_4_4 | [-21,-35,68]<br>[20,-33,69] |

*Continued...*

| Regions | Gyrus/Sulcus | Subregion Anatomical Description | Left/Right Hemisphere | ROI Label* | MNI Coordinates |
|---|---|---|---|---|---|
| Insular lobe | Insular gyrus (INS) | G, hypergranular insula | left<br>right | INS_L_6_1<br>INS_R_6_1 | [-36,-20,10]<br>[37,-18,8] |
| | | vIa, ventral agranular insula | left<br>right | INS_L_6_2<br>INS_R_6_2 | [-32,14,-13]<br>[33,14,-13] |
| | | dIa, dorsal agranular insula | left<br>right | INS_L_6_3<br>INS_R_6_3 | [-34,18,1]<br>[36,18,1] |
| | | vId/vIg, ventral dysgranular and granular insula | left<br>right | INS_L_6_4<br>INS_R_6_4 | [-38,-4,-9]<br>[39,-2,-9] |
| | | dIg, dorsal granular insula | left<br>right | INS_L_6_5<br>INS_R_6_5 | [-38,-8,8]<br>[39,-7,8] |
| | | dId, dorsal dysgranular insula | left<br>right | INS_L_6_6<br>INS_R_6_6 | [-38,5,5]<br>[38,5,5] |
| Limbic lobe | Cingulate gyrus (CG) | A23d, dorsal area 23 | left<br>right | CG_L_7_1<br>CG_R_7_1 | [-4,-39,31]<br>[4,-37,32] |
| | | A24rv, rostroventral area 24 | left<br>right | CG_L_7_2<br>CG_R_7_2 | [-3,8,25]<br>[5,22,12] |
| | | A32p, pregenual area 32 | left<br>right | CG_L_7_3<br>CG_R_7_3 | [-6,34,21]<br>[5,28,27] |
| | | A23v, ventral area 23 | left<br>right | CG_L_7_4<br>CG_R_7_4 | [-8,-47,10]<br>[9,-44,11] |
| | | A24cd, caudodorsal area 24 | left<br>right | CG_L_7_5<br>CG_R_7_5 | [-5,7,37]<br>[4,6,38] |
| | | A23c, caudal area 23 | left<br>right | CG_L_7_6<br>CG_R_7_6 | [-7,-23,41]<br>[6,-20,40] |
| | | A32sg, subgenual area 32 | left<br>right | CG_L_7_7<br>CG_R_7_7 | [-4,39,-2]<br>[5,41,6] |
| Occipital lobe | Medioventral occipital cortex (MVOcC) | cLinG, caudal lingual gyrus | left<br>right | MVOcC_L_5_1<br>MVOcC_R_5_1 | [-11,-82,-11]<br>[10,-85,-9] |
| | | rCunG, rostral cuneus gyrus | left<br>right | MVOcC_L_5_2<br>MVOcC_R_5_2 | [-5,-81,10]<br>[7,-76,11] |
| | | cCunG, caudal cuneus gyrus | left<br>right | MVOcC_L_5_3<br>MVOcC_R_5_3 | [-6,-94,1]<br>[8,-90,12] |
| | | rLinG, rostral lingual gyrus | left<br>right | MVOcC_L_5_4<br>MVOcC_R_5_4 | [-17,-60,-6]<br>[18,-60,-7] |
| | | vmPOS,ventromedial parietooccipital sulcus | left<br>right | MVOcC_L_5_5<br>MVOcC_R_5_5 | [-13,-68,12]<br>[15,-63,12] |
| | Lateral occipital cortex (LOcC) | mOccG, middle occipital gyrus | left<br>right | LOcC_L_4_1<br>LOcC_R_4_1 | [-31,-89,11]<br>[34,-86,11] |
| | | V5/MT+, area V5/MT+ | left<br>right | LOcC_L_4_2<br>LOcC_R_4_2 | [-46,-74,3]<br>[48,-70,-1] |
| | | OPC, occipital polar cortex | left<br>right | LOcC_L_4_3<br>LOcC_R_4_3 | [-18,-99,2]<br>[22,-97,4] |
| | | iOccG, inferior occipital gyrus | left<br>right | LOcC_L_4_4<br>LOcC_R_4_4 | [-30,-88,-12]<br>[32,-85,-12] |
| | | msOccG, medial superior occipital gyrus | left<br>right | LOcC_L_2_1<br>LOcC_R_2_1 | [-11,-88,31]<br>[16,-85,34] |
| | | lsOccG, lateral superior occipital gyrus | left<br>right | LOcC_L_2_2<br>LOcC_R_2_2 | [-22,-77,36]<br>[29,-75,36] |
| Subcortical nuclei | Thalamus (Tha) | Thalamus | left<br>right | Tha_L<br>Tha_R | [-12,-19,5]<br>[11,-18,6] |

*ROIs are labeled according to the Brainnetome atlas, except for posterior STS and thalamus where the left/right hemispheric masses are merged into a single ROI each.

**Supplementary Table 3. Twenty-nine active seed regions of interest (ROIs) identified using Pure /ta/ condition of all the 38 participants.**

| | Included subregions (mapped using Brainnetome atlas) | |
|---|---|---|
| **ROI name** | **LH seed** | **RH seed** |
| Superior frontal gyrus (SFG) | *medial areas 6,8,9*<br>*dorsolateral area 6* | *medial areas 6,8,9*<br>*lateral area 9* |
| Middle frontal gyrus (MFG) | *dorsal and ventral areas 9/46*<br>*IFJ; area 46* | *dorsal and ventral areas 9/46; IFJ; area 46*<br>*ventrolateral areas 6,8* |
| Inferior frontal gyrus (IFG) | *area 44; rostral area 45* | *area 44, IFS; caudal area 45* |
| Orbital gyrus (OrG) | *area 12/47* | *area 12/47* |
| Precentral gyrus (PrG) | *caudolateral area 6; area 4 (tongue/larynx region)*<br>*area 4 (head/face region)* | *caudolateral area 6*<br>*area 4 (tongue/larynx region)* |
| Superior temporal gyrus (STG) | *TE1.0 and TE1.2; area 41/42*<br>*rostral and caudal area 22* | *TE1.0 and TE1.2; lateral area 38*<br>*rostral and caudal area 22* |
| Middle temporal gyrus (MTG) | *aSTS, dorsolateral area 37* | *No activation* |
| Fusiform gyrus (FuG) | *medioventral area 37* | *No activation* |
| Posterior superior temporal sulcus (pSTS) | Posterior superior temporal sulcus | Posterior superior temporal sulcus |
| Superior parietal lobule (SPL) | *No activation* | *intraparietal area 7* |
| Inferior parietal lobule (IPL) | *caudal, rostroventral area 40*<br>*rostrodorsal areas 39, 40* | *caudal area 40; rostroventral area 40*<br>*rostrodorsal areas 39, 40* |
| Postcentral gyrus (PoG) | *area 1/2/3 (tongue/larynx)* | *No activation* |
| Insular gyrus (INS) | *dorsal and ventral agranular insula*<br>*dorsal granular and dysgranular insula* | *ventral agranular insula*<br>*dorsal granular and dysgranular insula* |
| Anterior cingulate cortex (ACC) | *No activation* | *pregenual area 32* |
| Medioventral occipital cortex (MVOcC) | *lingual and cuneus gyri* | *lingual and cuneus gyri* |
| Lateral occipital cortex (LOcC) | *inferior occipital gyrus; occipital polar cortex* | *inferior occipital gyrus* |
| Thalamus (Tha) | *left hemispheric combined thalamus* | *right hemispheric combined thalamus* |

* IFJ: inferior frontal junction; IFS: inferior frontal sulcus; aSTS: anterior superior temporal sulcus.

**Supplementary Table 4. Correlation between seed functional connectivity (whole brain) and the behavioral /ta/ response.**

| | High perceivers | | Low perceivers | |
|---|---|---|---|---|
| **Seed region** | **LH Mean corr.*** | **RH Mean corr.*** | **LH Mean corr.*** | **RH Mean corr.** |
| SFG | -0.39 | *ns* | 0.167 | -0.334 |
| MFG | 0.265 | *ns* | 0.442 | 0.438 |
| IFG | 0.352 | 0.161 | 0.463 | *ns* |
| OrG | 0.208 | 0.156 | -0.463 | *ns* |
| PrG | -0.222 | 0.109 | 0.313 | *ns* |
| STG | -0.433 | 0.463 | 0.118 | 0.12 |
| MTG | *ns* | NA | *ns* | NA |
| FuG | -0.368 | NA | 0.491 | NA |
| pSTS | 0.069 | 0.501 | 0.559 | 0.447 |
| SPL | NA | -0.511 | NA | 0.515 |
| IPL | -0.354 | *ns* | 0.183 | *ns* |
| PoG | *ns* | NA | 0.507 | NA |
| INS | -0.64 | 0.071 | 0.15 | 0.182 |
| ACC | NA | 0.381 | NA | 0.348 |
| MVOcC | -0.597 | -0.656 | 0.828 | 0.801 |
| LOcC | *ns* | *ns* | 0.285 | 0.807 |
| Tha | -0.696 | *ns* | 0.339 | 0.513 |

$^{*}p < 0.05$, FDR corrected; LH: left hemisphere; RH: right hemisphere; ns: non-significant.

### Supplementary Methods

The local clustering coefficient of a node estimates the fraction of connections among its first neighbors against all possible neighborhood connections, and the normalized averaged clustering coefficient ($C_n$) is defined as

$$C_n = \frac{C_{\text{real}}}{C_{\text{rand}}} \quad [1]$$

where $C_{\text{real}}$ and $C_{\text{rand}}$ are the average local clustering coefficients (1, 2) of the empirical and randomized networks, respectively.

Similarly, for the characteristics path length that measures the average shortest path across all node pairs, the normalized characteristics path length is

$$L_n = \frac{L_{\text{real}}}{L_{\text{rand}}} \quad [2]$$

where $L_{\text{real}}$ and $L_{\text{rand}}$ represent the characteristics path length (1) of the empirical and randomized networks, respectively.

Modularity on the other hand is a quality function that measures how well a network is divided into distinct functional units, and it can be conceptualized as another network segregation measure at the global scale (3). The normalized modularity is further estimated as (4)

$$Q_n = \frac{Q_{\text{real}} - Q_{\text{rand}}}{Q_{\text{max}} - Q_{\text{rand}}} \quad [3]$$

Here $Q_{\text{max}}$ is the maximal possible network modularity value with the same degree sequence and can be approximated as $Q_{\text{max}} = 1 - 1/m$, where $m$ represents the total number of modules in the real network (5). $Q_{\text{real}}$ and $Q_{\text{rand}}$ are modularity computed in the empirical and randomized network (3) respectively.

Node participation coefficient ($P$) measures how equally the connections of a node are distributed among all available modules of a network (6). Thus, it estimates the capacity of a node in intermodular information integration.

$$P = 1 - \sum_{i=1}^{m} \left( \frac{k_i}{k} \right)^2 \quad [4]$$

where $k$ is the degree of the node and $k_i$ represents the number of connections the node have with the module $i$. Here, $m$ is the number of available modules.
